# EZH2 Serine 21 Phosphorylation Restrains Compact-State PRC2 Activation and H3K27me3 Propagation

**DOI:** 10.64898/2026.06.02.729660

**Authors:** Andrew Q. Rashoff, Mark Matyas, Elizabeth G. Porter, Ryen Hazzard, Benjamin A. Garcia, Vignesh Kasinath, Peter W. Lewis

## Abstract

Polycomb Repressive Complex 2 (PRC2) propagates H3K27me3 through EED-dependent allosteric activation, yet how cells modulate the magnitude of this positive-feedback response remains poorly understood. Here, we identify phosphorylation of EZH2 serine 21 as a post-translational mechanism that attenuates PRC2 allosteric responsiveness. Prior structural studies have established that activator-bound PRC2 adopts both compact and extended active conformations. Using cryo-EM classification of wild-type and phospho-null EZH2 S21A PRC2 complexes, we find that the phospho-null EZH2 S21A substitution changes the distribution of particles across these pre-existing states, shifting PRC2 from a predominantly extended conformation to one enriched for the compact, allosterically activated conformation. Consistent with this structural transition, EZH2 S21A increases basal PRC2 activity, lowers the EC_50_ for H3K27me3-dependent stimulation, and accelerates H3K27me3 accumulation on peptide and nucleosome substrates. Disruption of the EED-EZH2 interface suppresses the S21A gain-of-activity phenotype, indicating that S21 phosphorylation constrains PRC2 by limiting productive EED-EZH2 allosteric coupling. In mesenchymal progenitor cells, loss of this phosphorylation-dependent restraint broadens H3K27me3 domains, reduces canonical PRC1 enrichment at high-occupancy Polycomb target loci, misregulates lineage-associated transcriptional programs, and impairs differentiation. These findings identify EZH2 S21 phosphorylation as a molecular rheostat that limits compact-state PRC2 activation, constrains H3K27me3 spreading, and preserves Polycomb-dependent developmental competence.

## Introduction

Development and tissue homeostasis require chromatin-based mechanisms that preserve gene-expression programs across cell divisions while permitting regulated transitions in cell identity. Polycomb group proteins were identified genetically as mediators of heritable developmental gene repression, and biochemical studies subsequently defined Polycomb Repressive Complex 2 (PRC2) as a conserved histone methyltransferase that catalyzes mono-, di-, and trimethylation of histone H3 lysine 27 (H3K27me1/2/3)^1–4^. The PRC2 core contains EZH1 or EZH2, EED, SUZ12, and RBBP4 or RBBP7. Accessory subunits assemble with this core to generate PRC2.1 and PRC2.2 complexes with distinct chromatin-binding and regulatory properties. PRC2.1 contains one Polycomb-like (PCL) family subunit together with EPOP or PALI1; these accessory subunits promote targeting to unmethylated CpG-rich Polycomb chromatin and tune PRC2 catalytic output through DNA- and chromatin-dependent mechanisms^5–10^. PRC2.2 contains AEBP2 and JARID2, which promote recognition of PRC1-dependent H2AK119ub and regulate PRC2 activity on nucleosomal substrates^8,11–14^. Thus, PRC2 output reflects both chromatin recruitment and the catalytic state of the enzyme after recruitment.

PRC2-mediated repression depends on mechanisms that promote H3K27 methylation at appropriate loci while limiting methylation in active chromatin. Positive inputs, including CpG island recognition, PRC1-mediated H2AK119ub, low transcriptional activity, and favorable nucleosomal substrate features, support PRC2 recruitment and activity at Polycomb target genes^15–18^. Conversely, active histone modifications including H3K4me3, H3K36me2/3, and RNA antagonize PRC2 activity and restrict the spread of H3K27 methylation into transcriptionally active regions^19–29^. Together, these opposing mechanisms focus PRC2 activity at repressed developmental genes while protecting active chromatin from inappropriate Polycomb-mediated silencing.

After recruitment and exclusion mechanisms define where PRC2 can act, the extent of local H3K27 methylation is further shaped by product-dependent allosteric activation. Binding of H3K27me3 to the EED aromatic cage stimulates the EZH2 SET domain, creating a read-write mechanism that reinforces H3K27me3 deposition and promotes propagation of Polycomb domains from nucleation sites into surrounding chromatin^30–32^. Related EED-dependent inputs are provided by methylated JARID2 in PRC2.2 and methylated PALI1 in PRC2.1, suggesting that histone-derived and cofactor-derived methyl-lysine ligands can engage a shared allosteric activation^12,33^. Because this positive feedback mechanism can amplify local H3K27 methylation, cells must tune the strength of allosteric activation to preserve domain boundaries and control of developmental genes.

Structural studies have focused on how trimethyl-lysine recognition by EED is transmitted to the EZH2 catalytic lobe. Allosteric ligand binding is coupled to rearrangements in EZH2 regulatory elements that include SANT1, the SANT1-binding domain (SBD), the stimulatory-responsive motif (SRM), and the SET activation loop (SAL)^34–36^. Cryo-electron microscopy (cryo-EM) analyses have identified two active conformations of human PRC2, termed extended active and compact active PRC2^14,37,38^. In the extended active state of PRC2, the SBD is straight, displacing SANT1 from the core, and SRM density is weak or absent due to its conformational flexibility^37,38^. By contrast, compact active PRC2 is considered the more catalytically competent state: the SBD adopts a bent configuration, positioning SANT1 against the catalytic core and allowing the ordering of the SRM, thereby facilitating the allosteric pathway linking EED to the EZH2 SET catalytic site. Together, these structures define the EZH2 regulatory module as the conformational link between EED ligand recognition, compact active-state formation, and catalytic activation.

Despite these structural and biochemical insights on PRC2 activation, how regulatory signals fine-tune PRC2 allostery remains poorly understood. Post-translational modifications on PRC2 subunits provide one potential mechanism for linking signaling pathways to PRC2 conformational state and catalytic output. Phosphorylation, methylation, and automethylation events on PRC2 components have been catalogued and linked to changes in chromatin binding, RNA engagement, stability, and enzymatic activity^39–43^. Among these modifications, phosphorylation of EZH2 serine 21 (S21-phos) has been associated with reduced H3K27me3, altered gene repression, and disease-associated signaling downstream of AKT, whereas phosphorylation at other EZH2 residues, including T311, attenuates PRC2 through distinct mechanisms^44–46^. However, how S21 phosphorylation influences the PRC2 allosteric cycle, the structural rearrangements of the enzyme complex, and the domain-scale distribution of H3K27me3 has remained unresolved. Addressing this question is necessary to understand how altered PRC2 activity can either reinforce aberrant repression or erode Polycomb-dependent chromatin states. These defects are hallmarks of B-cell lymphomas with EZH2 gain-of-function mutations and pediatric brain tumors where H3 K27M and EZHIP inhibit PRC2 activity to remodel the H3K27 methylation landscape^41,47–50^.

Here, we define EZH2 S21-phos as a post-translational input that tunes PRC2 allosteric responsiveness. We show that the phospho-null EZH2 S21A substitution increases basal activity and enhances responsiveness to H3K27me3-derived allosteric activator peptide. Using cryo-electron microscopy (cryo-EM), we find that loss of the S21-phos site enriches for the compact active state of PRC2, allowing us to determine a 3.4 Å structure of PRC2-S21A in the compact active state, suggesting a structural basis for the enhanced responsiveness to allosteric activation. In cells, EZH2 S21A broadens H3K27me3 domains without a comparable increase in EZH2 occupancy, reduces canonical PRC1 enrichment at Polycomb target loci, misregulates developmental gene expression programs, and impairs adipogenic and chondrogenic differentiation. Together, these findings define EZH2 S21 phosphorylation as a signaling-sensitive rheostat that limits PRC2 allosteric amplification, constrains H3K27me3 spreading, and preserves Polycomb domain fidelity during cell-fate control.

## Results

### PRC2 phosphorylation has a net negative effect on catalytic activity *in vitro*

Phosphorylation of PRC2 subunits has been extensively catalogued, with sites distributed across EZH2, SUZ12, EED, and RBBP4 and linked to changes in RNA binding, protein stability, chromatin association, and enzymatic output^39,40,42^. However, how the phosphorylation state of the intact complex affects PRC2 catalytic activity remains incompletely defined. To assess the net impact of phosphorylation on PRC2 methyltransferase activity, we compared purified recombinant human PRC2 before and after lambda phosphatase treatment using nucleosome-based methyltransferase assays. Phosphatase-treated PRC2 accumulated higher levels of H3K27me2/3 than untreated PRC2, indicating that phosphorylation constrains PRC2 catalytic output *in vitro* (**Figure 1A**). This increase was maintained across a titration of stimulatory H3K27me3 peptide, with phosphatase-treated PRC2 achieving higher H3K27 methylation at lower peptide concentrations than untreated PRC2. Orthogonal scintillation assays using ^3^H-SAM confirmed that phosphatase-treated PRC2 incorporated methyl groups more efficiently than untreated PRC2 (**Figure 1B**). These findings extend prior work showing that phosphorylation can suppress PRC2 methyltransferase activity^44,45^. Together, these data indicate that phosphorylation of recombinant PRC2 limits overall methyltransferase activity and attenuates the response to stimulation by the H3K27me3 peptide.

**Figure 1.**
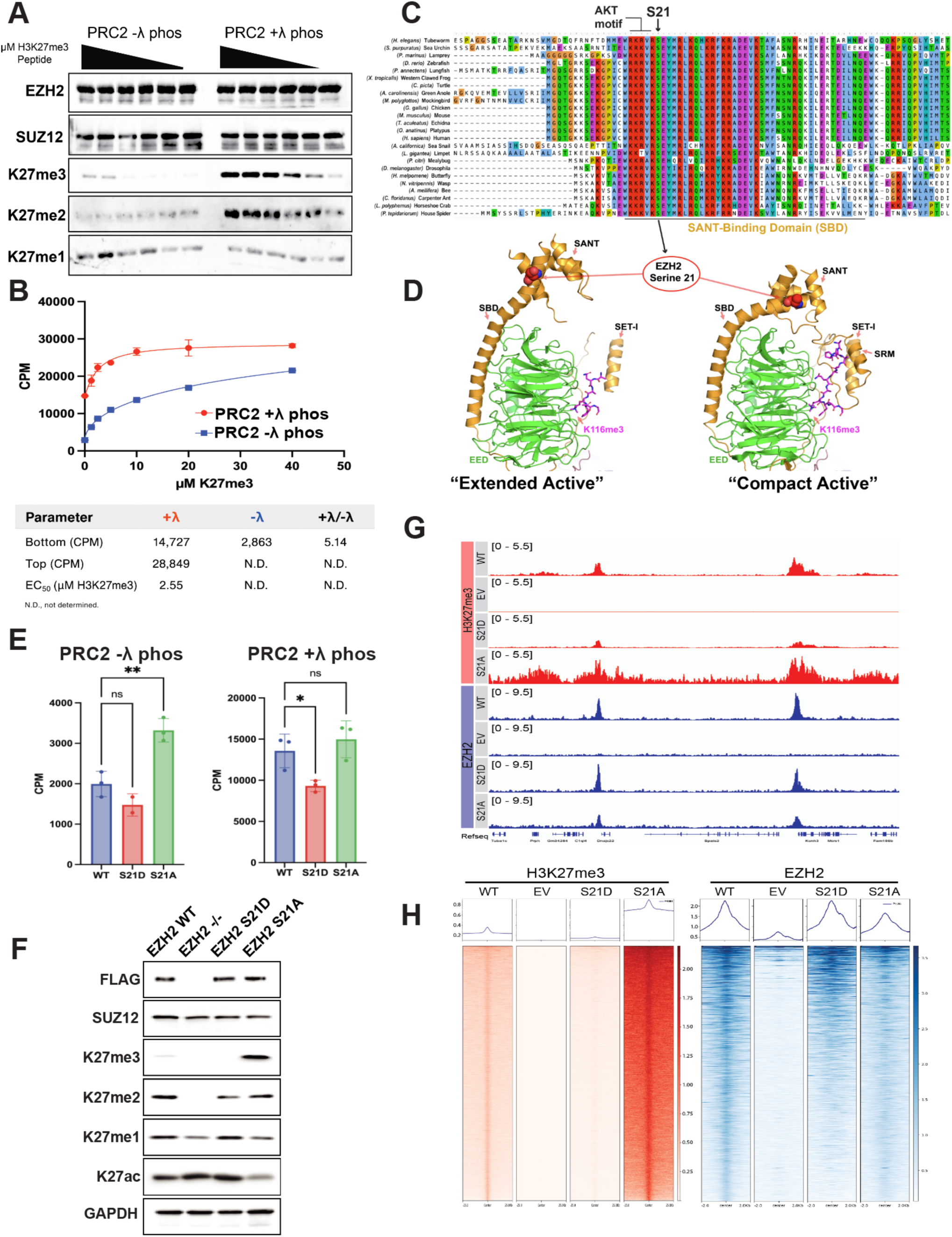
PRC2 phosphorylation reduces catalytic activity and attenuates H3K27 methylation. **A.** *In vitro* histone methyltransferase assays comparing untreated recombinant PRC2 and complex treated with lambda phosphatase. Sf9-purified PRC2 was incubated with recombinant nucleosomes and a 2-fold serial dilution of stimulatory H3K27me3 peptide beginning at 40 μM. **B.** Quantification of PRC2 activity in response to increasing concentrations of stimulatory H3K27me3 peptide. Untreated or lambda phosphatase-treated recombinant PRC2 was incubated with recombinant nucleosomes, and methyltransferase activity was measured by incorporation of ³H-SAM. Data are shown as mean ± SEM. Dose-response fit parameters are shown below the graph. **C.** Multiple sequence alignment of EZH2/EZH homologs from animal species showing conservation of the S21-containing AKT phosphorylation motif. Serine 21 lies within the SANT1-binding domain (SBD); the N-terminal segment of the SBD encompassing S21 is highly conserved across animals relative to the more divergent C-terminal segment. **D.** Structural localization of EZH2 serine 21 within the extended active and compact active PRC2 conformations (PDB 6C24 and 6C23, respectively). Serine 21 is positioned on the SBD near the junction with the SANT1 domain. The SBD-SANT1 region undergoes a large conformational rearrangement between the two states, with compact active PRC2 displaying a bent SBD and repositioned SANT1 domain. **E.** *In vitro* methyltransferase activity of recombinant PRC2 complexes containing wild-type EZH2, S21D, or S21A, assayed before or after lambda phosphatase treatment. Complexes were incubated with recombinant nucleosomes, and activity was quantified by incorporation of ³H-SAM. Data are shown as mean ± SEM, with individual replicate values indicated. Statistical comparisons are indicated above the bars. **F.** Immunoblot analysis of *Ezh2*-knockout MEFs reconstituted with FLAG-tagged wild-type EZH2, S21D, S21A, or empty vector. GAPDH serves as a loading control. **G.** Representative genome browser tracks showing spike-in-normalized H3K27me3 and EZH2 ChIP-seq signal at Polycomb target loci in *Ezh2*-knockout MEFs expressing the indicated EZH2 transgenes or empty vector. ChIP-seq experiments were normalized using exogenous spike-in chromatin. **H.** Heatmaps and aggregate profiles showing spike-in-normalized H3K27me3 and EZH2 ChIP-seq signal centered on the corresponding H3K27me3 or EZH2 peaks in *Ezh2*-knockout MEFs expressing the indicated EZH2 transgenes or empty vector. ChIP-seq experiments were normalized using exogenous spike-in chromatin.

### EZH2 serine 21 phosphorylation reduces PRC2 catalytic activity *in vitro* and in cells

Among PRC2-associated phosphorylation sites, EZH2 serine 21 emerged as a conserved modification previously implicated in negative regulation of H3K27 methylation, but the mechanism by which this phosphorylation event restrains PRC2 activity has remained unclear^42,44,45,51^. Alignment of EZH2/EZH homologs showed that the S21-containing AKT motif is highly conserved across animals and lies within the N-terminal segment of the SBD, whereas the C-terminal portion of the SBD is comparatively divergent (**Figure 1C**). Structural mapping placed S21 at the SBD-SANT1 junction, a conformationally flexible region that differs markedly between the extended active and compact active conformations (**Figure 1D**)^37,38^. In the extended active conformation, the SBD is comparatively straight and SANT1 is displaced from the catalytic core, whereas in the compact active conformation the SBD bends, SANT1 is repositioned against the core, and the SRM becomes ordered. This placement suggested that S21-phos could modulate PRC2 activity by tuning regulatory-module flexibility and EED-to-EZH2 allosteric communication rather than by altering complex abundance or recruitment. We therefore tested whether S21 contributes to the phosphorylation-dependent inhibition of PRC2 activity inferred from lambda phosphatase treatment. To assess the specific contribution of S21-phos, we performed methyltransferase assays with recombinant PRC2 complexes containing wild-type EZH2, phospho-null EZH2 S21A, or phosphomimetic EZH2 S21D, before and after lambda phosphatase treatment (**Figure 1E**). In agreement with prior work identifying S21-phos as inhibitory, untreated PRC2 containing S21A displayed higher catalytic activity than untreated wild-type PRC2, whereas PRC2 containing S21D was less active^44^. Notably, however, the S21A substitution accounted for only part of the increase in activity observed after phosphatase treatment, indicating that additional phosphorylation events cooperate with S21 to restrain PRC2 catalytic output *in vitro* (**Figure 1E**).

We next tested the cellular consequences of perturbing this conserved S21 phosphorylation site by expressing wild-type EZH2, S21A, S21D, or empty vector in *Ezh2*-knockout mouse embryonic fibroblasts and assessed global H3K27 methylation. Immunoblotting of K27 methylation and acetylation revealed a pronounced difference between the cell lines with different EZH2 constructs (**Figure 1F**). Expression of the phospho-null S21A mutant resulted in an increase in global H3K27me3, consistent with a negative regulatory role for S21-phos across multiple cellular contexts^44,51^. Expression of EZH2 S21A also reduced H3K27ac, consistent with redistribution of H3K27 from acetylated toward methylated states in S21A-expressing cells (**Figure 1F**).

To determine whether these effects reflected altered PRC2 composition rather than a direct change in catalytic regulation, we asked whether S21-phos alters PRC2 assembly or association with accessory subunits that define PRC2.1 and PRC2.2 complexes^8^. FLAG immunoprecipitation from nuclear extracts of *Ezh2*-knockout MEFs expressing wild-type EZH2, S21A, S21D, or empty vector showed comparable co-recovery of core PRC2 components and representative accessory subunits, without evident genotype-dependent differences (**Figure S1A**). These results indicate that the increased H3K27 methylation observed with EZH2 S21A is unlikely to arise from altered PRC2 complex composition and instead supports a direct role for S21-phos in restraining PRC2 enzymology and H3K27me3 deposition.

To quantify these changes and place them in the context of other histone modifications, we performed quantitative mass spectrometry on acid-extracted histones from the same MEF lines. This analysis revealed broad changes in H3 post-translational modification states in cells expressing S21A or S21D relative to wild-type EZH2 (**Figure S1B**). Relative H3K27 methylation stoichiometries were altered in all transgene conditions compared with wild-type EZH2, with S21A expression increasing H3K27me3 and decreasing H3K27me2, H3K27me1, and unmodified H3K27 (**Figure S1C**). Relative ratios of H3K36 methylation states were also altered in S21A-expressing cells compared with wild type (**Figure S1D**). Because H3K36 methylation antagonizes PRC2 activity and limits H3K27me3 spreading from nucleation sites, we examined H3K27 methylation on peptides carrying H3K36me1, H3K36me2, or H3K36me3^19,20,25,52^. EZH2 S21A increased the fraction of H3K27-methylated species detected among H3K36-methylated peptides, indicating that the steady-state increase in H3K27 methylation extends to histones bearing H3K36 methylation (**Figure S1E-G**).

To assess how S21-phos shapes chromatin-level outputs of PRC2, we performed ChIP-seq for H3K27me3 and EZH2 in *Ezh2*-knockout MEFs expressing wild type, S21A, S21D, or empty vector. As expected from the immunoblot and mass spectrometry data, S21A expression produced a global increase in H3K27me3, whereas S21D expression decreased H3K27me3 relative to wild-type (**Figure 1G, 1H**). Although H3K27me3 is markedly reduced by immunoblotting in EZH2 S21D-expressing cells, ChIP-seq detects residual H3K27me3 at Polycomb target regions, indicating that S21D attenuates, but does not abolish, PRC2-dependent H3K27me3 deposition. In S21A-expressing cells, H3K27me3 signal increased both at sites of initial PRC2 recruitment and at previously defined “spread” regions associated with allosterically activated PRC2, and in many cases extended into genomic domains that lacked appreciable H3K27me3 in wild-type cells (**Figure 1G, 1H**)^32,49^. By contrast, EZH2 occupancy at recruitment sites was broadly similar among wild-type, S21A, and S21D conditions, indicating that the marked differences in H3K27me3 arise primarily from altered catalytic output rather than broad changes in chromatin binding. This finding contrasts with earlier models in which loss of S21-phos was proposed to enhance PRC2 chromatin association^44^ and instead supports a mechanism in which S21-phos directly tunes PRC2 catalytic responsiveness to allosteric inputs.

### EZH2 S21 phosphorylation limits PRC2 occupancy of the compact active conformation

Because S21A increases PRC2 catalytic output without broadly altering EZH2 chromatin occupancy or PRC2 complex composition, we asked whether S21-phos regulates the conformational equilibrium of the complex itself. Previous structural work on PRC2 showed that Serine 21 lies within the EZH2 N-terminal regulatory region, near the SBD-SANT1-SRM module that distinguishes compact active and extended active PRC2 conformations (**Figure 1D**). We therefore used cryo-EM to compare recombinant PRC2 complexes containing either wild-type EZH2 or phospho-null EZH2 S21A, together with SUZ12, EED, RBBP4, AEBP2, and JARID2(aa 106-350) (**Figure 2A**). Previous structural and biochemical work has shown that a co-expressed N-terminal fragment of JARID2 (aa 106-350) containing trimethylated K116 is sufficient to bind both EED and SUZ12 and serves as an allosteric activator^37^. This places PRC2 in an activator-bound, or allosterically primed, state from which it can sample extended active and compact active conformations. To facilitate an unbiased comparison of the conformational differences between PRC2 containing either wild-type EZH2 or phospho-null EZH2 S21A, protein purification, cryo-EM sample preparation, data collection, and data processing were carried out identically across both data samples (**Figure S2; Figure S3A, B, Table 1)**^37,38^. This led us to determine high resolution reconstructions of wild-type and S21A PRC2 in both active conformations.

**Figure 2.**
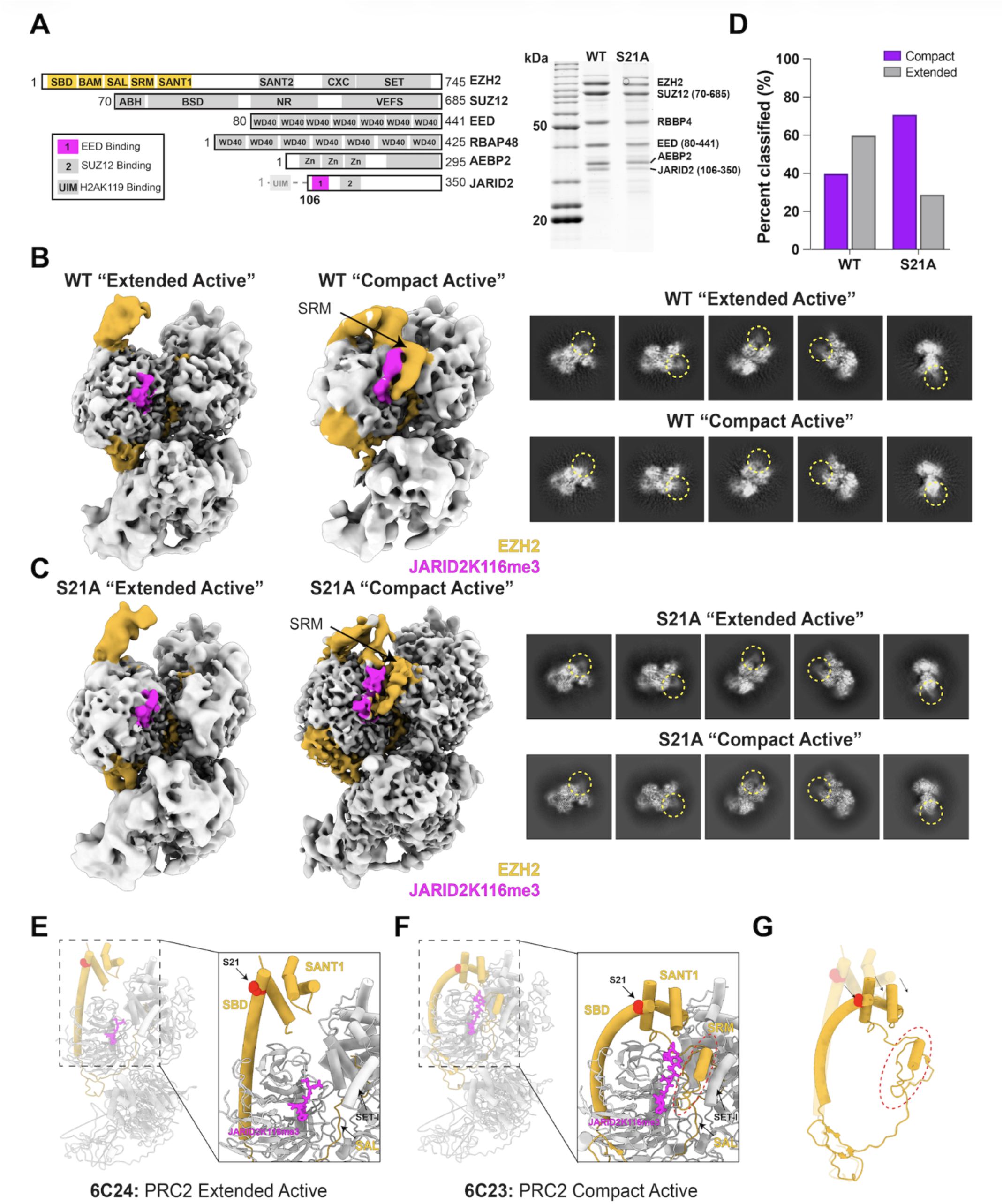
Activator-bound PRC2 samples extended and compact active conformations, and loss of EZH2 S21 phosphorylation enriches the compact active state. **A.** Domain schematic and Coomassie blue-stained SDS-PAGE analysis of recombinant human PRC2 used for cryo-EM. Complexes contained wild-type (WT) EZH2 or phospho-null EZH2 S21A together with SUZ12 (aa 70-685), EED (aa 80-441), RBBP4, AEBP2, and JARID2 (aa 106-350). The JARID2 fragment contains the K116me3 EED-binding motif and was included to provide an allosteric activating ligand; AEBP2 was included to stabilize PRC2 during cryo-EM analysis. **B, C.** Locally filtered cryo-EM maps of WT PRC2 (**B**) and EZH2 S21A PRC2 (**C**) in extended active and compact active conformations. The EZH2 regulatory module is shown in gold, and the JARID2 K116me3 EED-binding segment is shown in magenta. In the compact active, allosterically activated conformation of both WT and S21A PRC2, density for the stimulatory-responsive motif (SRM) is ordered and visible. Representative 2D class averages for the extended and compact classes are shown to the right; yellow dashed circles mark density corresponding to the rearranged SBD-SANT1 region. **D.** Distribution of classified particles after heterogeneous refinement of WT and S21A PRC2 datasets using 20 Å low-pass-filtered input maps and no forced hard classification. Particle distributions were WT compact, 40.3%; WT extended, 59.7%; S21A compact, 71.3%; and S21A extended, 28.7%. **E, F.** Previously determined PRC2 structures representing the extended active (PDB 6C24; **E**) and compact active (PDB 6C23; **F**) conformations. Insets highlight structural rearrangements that distinguish the two active states, including bending of the SBD helix, repositioning of SANT1, and ordering of the SRM in the compact active state. EZH2 S21 is shown as red spheres. **G.** Schematic summary of the S21 phosphorylation-sensitive rearrangement of the EZH2 regulatory module. Loss of S21 phosphorylation, modeled by phospho-null EZH2 S21A, increases the fraction of PRC2 particles in the compact active conformation, characterized by a bent SBD, SANT1 repositioning against the catalytic core, and SRM ordering.

Cryo-EM data processing revealed that wild-type and EZH2 S21A PRC2 sampled two major active conformational states, in agreement with previous structural studies of wild-type PRC2 (**Figure 2B, C**)^37^. The dominant structural consequence of S21A, however, was a striking, nearly reciprocal inversion of the active-state distribution. Wild-type PRC2 was predominantly extended active, with approximately 60% of classified particles in the extended state and 40% in the compact state. By contrast, EZH2 S21A was predominantly compact active, with approximately 71% of particles in the compact state and only 29% in the extended state (**Figure 2D).** The same qualitative shift was reproduced under alternative classification conditions **(Figure S3C, D).** Thus, loss of the S21-phos site switches the predominant PRC2 population from the extended active state to the compact, allosterically activated state. This near-reciprocal redistribution identifies S21-phos as a major determinant of the conformational equilibrium of activator-bound PRC2 and provides a structural basis for the enhanced allosteric responsiveness of S21A PRC2. The structural features that distinguish the two states are illustrated by the previously determined extended and compact active models (**Figure 2E, F**). In the extended active state, the SBD remains comparatively straight, SANT1 is displaced from the catalytic core, and SRM density is weak or absent (**Figure 2E**). In the compact active state, the SBD bends, SANT1 is repositioned against the catalytic core, and the SRM becomes ordered, completing the SANT1-SBD-SRM architecture associated with allosteric activation (**Figure 2F**)^34,35,37^. Despite this pronounced redistribution between active states, the overall PRC2 architecture remained closely similar across wild-type and S21A reconstructions (**Figure 2B, C**), and direct SET-domain fitting confirmed close agreement across the maps **(Figure S3E**). The compact- and extended-active SET domains also superimposed closely when aligned on either the SET domain or EED (**Figure S3F**), indicating that S21A shifts occupancy of a pre-existing active conformation rather than generating a distinct PRC2 architecture.

Together, these structural data suggest that EZH2 S21-phos restrains sampling of the compact, allosterically activated PRC2 conformation. By shifting the conformational equilibrium toward this state, the S21A substitution provides a structural basis for enhanced product-stimulated PRC2 activity. These findings place EZH2 S21-phos within the conformational control mechanism that links EED-dependent allosteric priming to compact-state activation and EZH2 catalytic output. This S21-phos-sensitive rearrangement is schematized in **Figure 2G**.

### EZH2 S21 phosphorylation modulates response to H3K27me3-mediated allosteric priming

Our cryo-EM analyses indicate that EZH2 S21A shifts PRC2 toward the compact, allosterically activated conformation. We next asked whether this conformational shift alters PRC2 biochemical activity by altering substrate engagement or responsiveness to H3K27me3-mediated allosteric priming. Prior structural models suggested that elements within or near the EZH2 SBD can contact nucleosomal DNA and the H3 tail, raising the possibility that S21-phos could influence nucleic acid or nucleosome binding^34,38^. To test this directly, we performed electrophoretic mobility shift assays with recombinant wild-type PRC2 and PRC2 containing S21A, using Cy5-labeled 601 DNA alone or Cy5-labeled 601 DNA assembled into nucleosomes. Across a broad range of PRC2 concentrations, wild-type and S21A complexes showed no detectable difference in apparent binding affinity for either DNA or nucleosomes, indicating that the S21A mutation does not measurably alter bulk nucleic acid or nucleosome binding under these conditions (**Figure S4A, B**).

We therefore asked whether S21A instead alters the response to H3K27me3-mediated allosteric priming. In methyltransferase assays using H3 peptide substrates and increasing concentrations of stimulatory H3K27me3 peptide, PRC2 containing S21A showed elevated basal activity and a lower EC_50_ for H3K27me3-dependent stimulation compared with wild-type PRC2 (**Figure 3A**). Thus, loss of S21-phos increases PRC2 responsiveness to the product-derived allosteric activator. To determine how S21A alters enzyme kinetics across graded allosteric stimulation, we measured initial velocities over a range of H3(18–37) peptide substrate concentrations in the presence of increasing concentrations of stimulatory H3K27me3 peptide and fit the data to the Michaelis-Menten equation (**Figure 3B**). The fitted kinetic parameters are summarized in **Figure S4C**.

**Figure 3.**
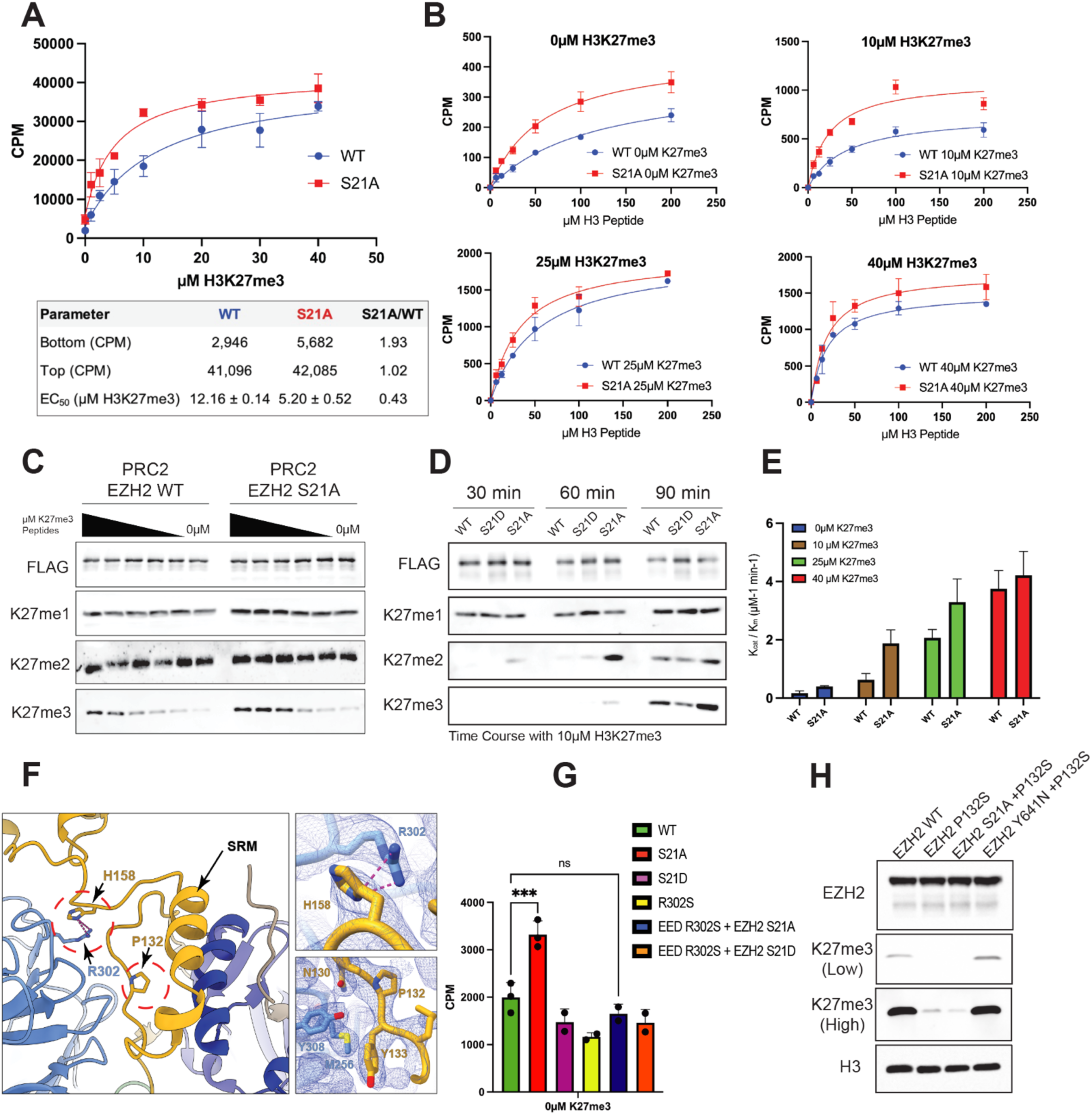
EZH2 S21 phosphorylation modulates PRC2 responsiveness to H3K27me3-dependent allosteric activation. **A.** *In vitro* methyltransferase assays comparing recombinant PRC2 complexes containing wild-type EZH2 or EZH2 S21A across increasing concentrations of stimulatory H3K27me3 peptide. Reactions were performed with ³H-SAM, and product formation was quantified by liquid scintillation counting of incorporated radiolabel. Data were fit by nonlinear regression using a three-parameter dose-response model. Fitted Bottom, Top, and EC50 values are shown in the table below the graph. Data are shown as mean ± SEM. **B.** Michaelis-Menten kinetic analysis of recombinant PRC2 complexes containing wild-type EZH2 or EZH2 S21A. Initial velocities were measured using histone H3(18-37) peptide substrate across the indicated substrate concentrations in the presence of 0, 10, 25, or 40 μM stimulatory H3K27me3 peptide. Reactions contained 20 nM PRC2, and initial velocities were quantified as CPM. Data were fit by nonlinear regression to the Michaelis-Menten equation and are shown as mean ± SD from three independent experiments. **C.** Immunoblot analysis of *in vitro* methyltransferase assays using recombinant PRC2 containing wild-type EZH2 or EZH2 S21A. Complexes were incubated with purified oligonucleosomes and SAM in the presence of a serial dilution of stimulatory H3K27me3 peptide, including a no-peptide control. **D.** Time-course analysis of *in vitro* methyltransferase activity for recombinant PRC2 complexes containing wild-type EZH2, EZH2 S21D, or EZH2 S21A. Complexes were incubated with purified oligonucleosomes and SAM in the presence of 10 μM stimulatory H3K27me3 peptide, and reactions were quenched at the indicated time points with SDS loading buffer. Reaction products were analyzed by immunoblotting. **E.** Catalytic efficiency of recombinant PRC2 containing wild-type EZH2 or EZH2 S21A in the presence of the indicated concentrations of stimulatory H3K27me3 peptide. Catalytic efficiency is plotted as kcat/Km and was derived from the kinetic fits shown in **B**. Data are shown as mean ± SEM. **F.** Structural view of the EED-EZH2 interface within the PRC2 allosteric coupling module (PDB 6WKR). The EZH2 SRM and residues implicated in allosteric signal transmission, including EED R302, EZH2 P132, and EZH2 H158, are indicated. Insets show the model docked into the experimentally determined EZH2 S21A PRC2 cryo-EM map (EMD-76889). Distances between histidine ND1 and arginine NH1/NH2 atoms are approximately 3-3.5 Å, consistent with a possible cation-π interaction between EZH2 H158 and EED R302. **G.** *In vitro* histone methyltransferase assays comparing recombinant PRC2 containing wild-type EZH2, EZH2 S21A, or EZH2 S21D in combination with wild-type EED or the allosteric coupling-deficient EED R302S mutant. Reactions contained 20 nM PRC2 and 20 μM H3 peptide substrate and were performed in the absence of stimulatory H3K27me3 peptide. Incorporation of ³H-methyl groups from ³H-SAM was quantified by liquid scintillation counting, and CPM values were normalized to no-substrate controls. Data are shown as mean ± SD, with individual replicate values indicated. Statistical comparisons are indicated above the bars. **H.** Immunoblot analysis of whole-cell lysates from 293T cells stably expressing wild-type EZH2, EZH2 P132S, EZH2 S21A/P132S, or EZH2 Y641N/P132S. EZH2 expression and cellular H3K27me3 were assessed by immunoblotting. Low and high exposures of the H3K27me3 immunoblot are shown, and total H3 serves as a loading control.

Consistent with these results, nucleosome methyltransferase assays analyzed by immunoblotting showed that S21A PRC2 accumulated H3K27me3 at lower concentrations of stimulatory H3K27me3 peptide than wild-type PRC2 (**Figure 3C**). Time-course assays on oligonucleosome substrates in the presence of 10 μM stimulatory H3K27me3 peptide further showed that S21A PRC2 accumulated H3K27me2 and H3K27me3 more rapidly than wild-type or S21D PRC2 (**Figure 3D**). At low concentrations of H3K27me3 peptide, S21A PRC2 exhibited higher catalytic efficiency than wild-type PRC2, as reflected by increased k_cat_/K_m_ (**Figure 3E**). At higher stimulator concentrations, the catalytic efficiencies of wild-type and S21A PRC2 converged, indicating that the S21A-dependent advantage is most evident under basal or subsaturating allosteric stimulation. Notably, H3K27me3-dependent stimulation of wild-type PRC2 increased k_cat_ and decreased K_m_ relative to the unstimulated state, indicating that product-derived allosteric activation enhances both turnover and apparent substrate engagement (**Figure S4C**).

Together, these biochemical data indicate that EZH2 S21-phos restrains PRC2 responsiveness to H3K27me3-mediated allosteric priming. Loss of the S21-phos site lowers the threshold for product-stimulated activation and accelerates H3K27me2/3 accumulation without measurably changing bulk DNA or nucleosome binding. Coupled with our cryo-EM results, these findings support a model in which EZH2 S21-phos limits access to the compact, allosterically activated PRC2 conformation, thereby attenuating catalytic amplification by H3K27me3.

### EZH2 S21A-enhanced PRC2 activity requires productive EED-EZH2 allosteric coupling

To test whether the enhanced activity of S21A-containing PRC2 requires productive EED-EZH2 allosteric coupling, we introduced S21 substitutions into PRC2 complexes carrying mutations that disrupt critical interactions between EED and the EZH2 catalytic lobe. Structural and biochemical studies identified critical residues at the EED-EZH2 allosteric interface for transmitting the H3K27me3-dependent allosteric priming signal from the EED aromatic cage to the EZH2 SET domain via the EZH2 (SRM)^34,49,53^. Of note, EED R302 contacts EZH2 H158 and anchors the SRM interface, whereas EZH2 P132 is positioned between the SRM and SET-I domains (**Figure 3F**). Our S21A compact active EM map agrees closely with these observations (**Figure 3F**), whereas these interactions are not observed in the extended active state.

We first tested this relationship *in vitro* using recombinant PRC2 complexes containing either wild-type EED or the allosteric activation-coupling-deficient EED R302S mutant, together with wild-type EZH2, S21A, or S21D. In the absence of exogenous stimulatory H3K27me3 peptide, S21A increased PRC2 activity relative to wild-type EZH2 when EED was intact (**Figure 3G**). This activity-enhancing effect was lost in complexes containing EED R302S, indicating that the elevated basal activity of S21A-containing PRC2 requires an intact EED-EZH2 allosteric interface (**Figure 3G**). Thus, S21A does not simply increase catalytic output independently of PRC2 regulatory state but instead acts through the EED-EZH2 allosteric communication pathway that mediates H3K27me3-dependent stimulation.

We next asked whether productive EED-EZH2 allosteric coupling is also required for the S21A-dependent increase in H3K27me3 in cells. We expressed wild-type EZH2, allosteric-defective EZH2 P132S, and the double mutant EZH2 S21A/P132S in 293T cells and monitored H3K27me3 by immunoblotting (**Figure 3H**). Consistent with prior work showing that P132S disrupts allosteric communication between EED and the EZH2 catalytic lobe, EZH2 P132S reduced H3K27me3 relative to wild-type EZH2^49,53^. Introduction of S21A into the P132S background did not restore H3K27me3, indicating that the S21A-dependent increase in cellular H3K27me3 requires an intact allosteric interface (**Figure 3H**).

As a comparison, we examined the EZH2 gain-of-function Y641N substitution, which increases H3K27me3 by altering substrate preference toward H3K27me2^54^. In contrast to S21A, the P132S/Y641N double mutant maintained H3K27me3 levels comparable to wild-type EZH2, consistent with a mechanism in which Y641N can increase H3K27me3 through altered catalytic specificity rather than the S21A-linked allosteric activation pathway (**Figure 3H**). Together, these *in vitro* and cellular data indicate that the enhanced activity of EZH2 S21A-containing PRC2 depends on canonical EED-EZH2 allosteric coupling and support a model in which S21-phos restrains catalytic output by limiting access to the compact, allosterically activated PRC2 conformation.

### EZH2 S21A disrupts Polycomb-dependent mesenchymal lineage control

Because EZH2 S21A increased PRC2 allosteric responsiveness and broadened H3K27me3 deposition, we next asked whether loss of this regulatory phosphorylation alters cell fate control in a differentiation-sensitive developmental context. Mesenchymal lineage commitment provides a tractable setting for testing how quantitative changes in histone methylation and Polycomb regulation are converted into durable changes in transcriptional output and developmental potential. Prior studies have used mesenchymal progenitor systems to show that oncohistone-driven disruption of H3K36 or H3 N-terminal residues can remodel H3K27me3 domains, alter PRC2-dependent gene regulation, and impair differentiation, underscoring the sensitivity of mesenchymal lineage programs to perturbations in chromatin state ^21,55–57^. We therefore tested whether enhanced PRC2 activity caused by EZH2 S21A perturbs this regulatory balance and compromises adipogenic and chondrogenic differentiation.

To test this possibility, we generated Ezh2-knockout murine mesenchymal progenitor cells and reconstituted them with wild-type EZH2, S21D, S21A, or an empty vector control. Immunoblot analysis confirmed loss of endogenous EZH2 and showed that S21A increased global H3K27me2/3 relative to wild-type EZH2 and S21D (Figure 4A). Thus, the S21A-dependent increase in PRC2 catalytic output is maintained in a mesenchymal progenitor context, providing a system to determine whether enhanced H3K27 methylation affects differentiation potential.

**Figure 4.**
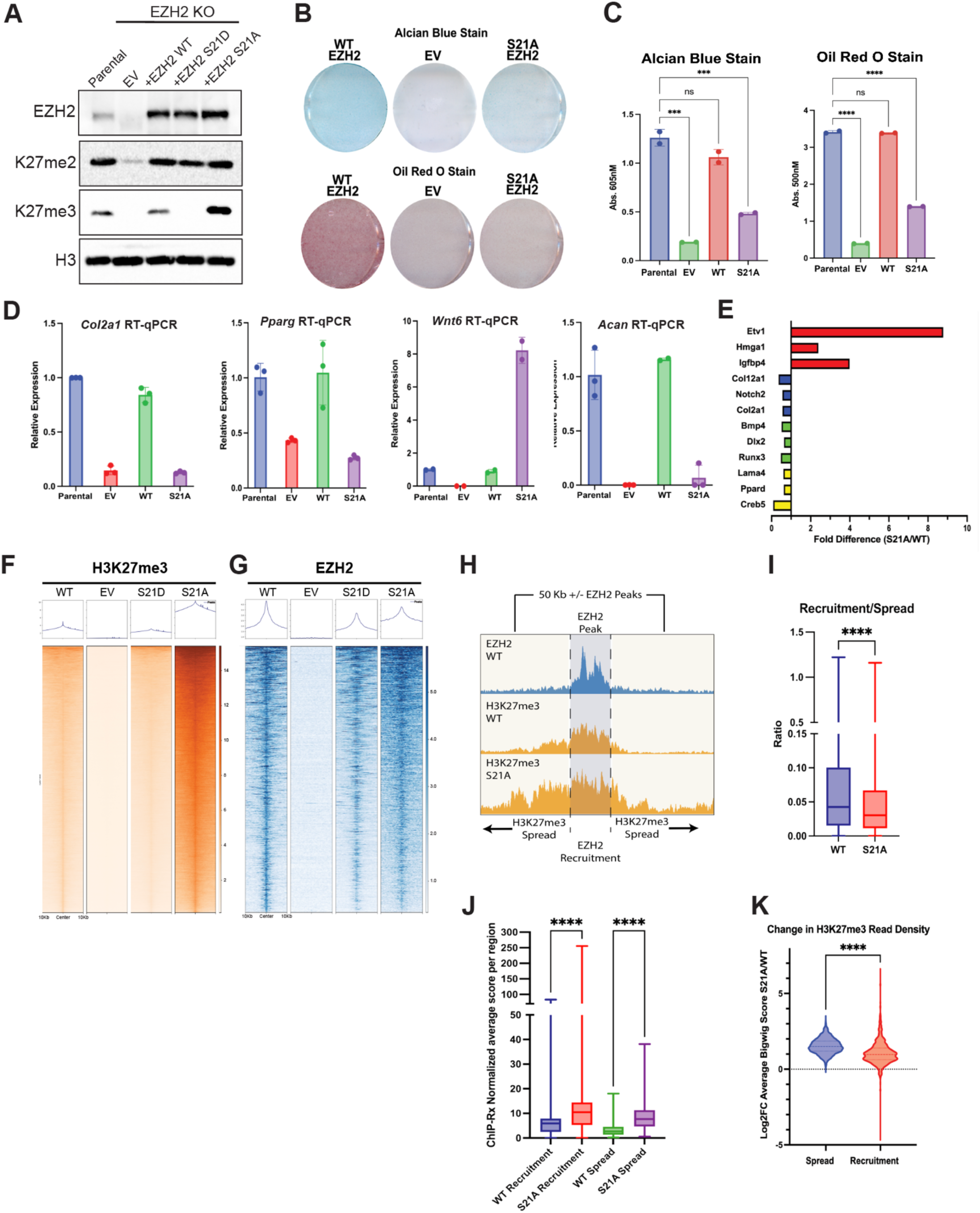
EZH2 S21A expands H3K27me3 domains and impairs mesenchymal differentiation. **A.** Immunoblot analysis of parental C3H10T1/2 mesenchymal progenitor cells and *Ezh2*-knockout cells reconstituted with empty vector, wild-type EZH2, EZH2 S21D, or EZH2 S21A. Blots were probed for EZH2, H3K27me2, H3K27me3, and total H3. **B.** Representative Alcian blue staining after 9 days under chondrogenic differentiation conditions (top) and Oil Red O staining after 9 days under adipogenic differentiation conditions (bottom) in *Ezh2*-knockout C3H10T1/2 cells expressing wild-type EZH2, empty vector, or EZH2 S21A. **C.** Quantification of Alcian blue and Oil Red O staining in parental cells and *Ezh2*-knockout cells expressing empty vector, wild-type EZH2, or EZH2 S21A. Alcian blue staining was solubilized and measured at 605 nm, and Oil Red O staining was extracted and measured at 500 nm. Bars represent mean ± SD, with individual replicate values indicated. Statistical comparisons are indicated above the bars. **D.** RT-qPCR analysis after 2 days under lineage-specific differentiation conditions. The chondrogenic markers *Col2a1* and *Acan* were measured under chondrogenic conditions, and the adipogenic differentiation-associated transcripts *Pparg* and *Wnt6* were measured under adipogenic conditions. Parental cells and *Ezh2*-knockout cells expressing empty vector, wild-type EZH2, or EZH2 S21A were analyzed. Bars represent mean ± SD from technical triplicates, with individual replicate values indicated. **E.** RNA-seq-derived fold differences in representative genes associated with mesenchymal stem cell proliferation (red), chondrocyte differentiation (blue), osteocyte differentiation (green), and adipocyte differentiation (yellow), comparing EZH2 S21A- and wild-type EZH2-expressing cells. The displayed genes are *Etv1*, *Hmga1*, *Igfbp4*, *Col12a1*, *Notch2*, *Col2a1*, *Bmp4*, *Dlx2*, *Runx3*, *Lama4*, *Ppard*, and *Creb5*. **F, G.** Heatmaps and aggregate profiles of ChIP-Rx-normalized H3K27me3 (**F**) and EZH2 (**G**) signal centered on the corresponding called peaks in *Ezh2*-knockout C3H10T1/2 cells expressing wild-type EZH2, empty vector, EZH2 S21D, or EZH2 S21A. Each row represents an individual peak ordered by signal intensity, and the plots span 10 kb on either side of the peak center. **H.** Representative profiles illustrating the analysis of H3K27me3 spreading from an EZH2 recruitment site. EZH2 signal defines the central recruitment interval, and H3K27me3 signal in wild-type EZH2- and EZH2 S21A-expressing cells is shown across the recruitment interval and the 50-kb flanking spread regions. **I.** Box-and-whisker plot showing the ratio of H3K27me3 ChIP-Rx signal at EZH2 recruitment sites to H3K27me3 signal within the flanking spread regions for each EZH2 peak, comparing wild-type EZH2 and EZH2 S21A. **J.** Box-and-whisker plots showing the normalized average H3K27me3 ChIP-Rx score per region at EZH2 recruitment sites and spread regions in wild-type EZH2- and EZH2 S21A-expressing cells. **K.** Violin plots showing the log2 fold change in H3K27me3 ChIP-Rx read density between EZH2 S21A- and wild-type EZH2-expressing cells at spread and recruitment regions. Statistical comparisons are indicated above the plots.

We next induced chondrogenic and adipogenic differentiation and assessed lineage output by histochemical staining and quantitative assays. Representative images after 9 days showed reduced Alcian blue staining under chondrogenic conditions and reduced Oil Red O staining under adipogenic conditions in *Ezh2*-knockout cells expressing empty vector or S21A relative to cells reconstituted with wild-type EZH2 (**Figure 4B**). Quantification confirmed impaired chondrogenic matrix deposition and lipid accumulation in these conditions (**Figure 4C**). These defects were also evident morphologically during early differentiation, as S21A-expressing and Ezh2-knockout cultures showed attenuated features of chondrocyte and adipocyte lineage commitment (**Figure S5C**).

Consistent with the staining phenotypes, RT-qPCR analysis showed impaired induction of lineage-associated transcriptional programs in S21A-expressing cells relative to cells reconstituted with wild-type EZH2. EZH2 S21A-expressing cells exhibited reduced expression of the chondrogenic markers *Col2a1* and *Acan* after chondrogenic induction^58^. After adipogenic induction, S21A-expressing cells showed reduced *Pparg* and increased *Wnt6* expression, consistent with disruption of the adipogenic gene expression program (**Figure 4D**)^59^. RNA-seq-derived fold differences further showed increased expression of representative transcripts associated with mesenchymal progenitor cell proliferation or maintenance and reduced expression of representative transcripts associated with chondrocyte, osteocyte, and adipocyte differentiation in S21A-expressing cells (**Figure 4E**). Together, these data indicate that loss of EZH2 S21-phos increases H3K27 methylation in mesenchymal progenitors and compromises their ability to execute adipogenic and chondrogenic differentiation programs.

### Enhanced H3K27me3 spreading in EZH2 S21A cells redistributes canonical PRC1

We next asked how the non-equivalent empty-vector and S21A differentiation phenotypes relate to canonical PRC1 localization. Empty-vector cells lack restored EZH2-dependent H3K27me3, whereas S21A cells retain abundant H3K27me3 that is distributed across broader domains. We therefore focused on how S21A-driven H3K27me3 spreading reshapes the relationship between PRC2 occupancy, H3K27 methylation, and canonical PRC1 localization in mesenchymal progenitor cells. Because canonical PRC1 complexes are recruited in part through CBX chromodomains that recognize H3K27me3, changes in the breadth, density, or distribution of H3K27me3 domains can reorganize canonical PRC1 occupancy at Polycomb target loci^21,60–63^. We first examined H3K27me3 and EZH2 occupancy in *Ezh2*-knockout cells expressing the different EZH2 transgenes. ChIP-Rx-normalized heatmaps and aggregate profiles showed that S21A increased H3K27me3 across Polycomb target regions, with comparatively modest changes in EZH2 localization (**Figure 4F, G**). ChIP-qPCR at representative PRC2 target loci and negative-control regions further supported increased H3K27me3 deposition in EZH2 S21A-expressing cells (**Figure S5A**).

To determine where the additional H3K27me3 accumulates, we partitioned EZH2-bound regions into recruitment sites and surrounding spread regions, defined as 50-kb windows flanking each EZH2 peak. Representative profiles illustrated the central EZH2 recruitment interval and the flanking regions used to quantify H3K27me3 spreading (**Figure 4H**). Across EZH2 peaks, S21A reduced the ratio of H3K27me3 signal at recruitment sites relative to flanking spread regions (**Figure 4I**). Separate quantification of normalized H3K27me3 signal at recruitment and spread regions showed increased deposition in S21A-expressing cells (**Figure 4J**), and region-specific fold-change analysis showed a greater S21A-dependent increase across spread regions than recruitment regions (**Figure 4K**). Thus, S21A shifts the H3K27me3 landscape from a more recruitment-focused pattern toward broader domains surrounding PRC2-bound loci.

We next examined how this remodeled H3K27me3 landscape affects canonical PRC1 localization. At representative Polycomb target loci, genome browser tracks showed reduced CBX2 and RING1B occupancy in EZH2 S21A-expressing cells despite increased H3K27me3 (**Figure 5A**). ChIP-qPCR at selected loci provided orthogonal support for reduced CBX2 and RING1B occupancy despite increased H3K27me3 (**Figure S5A**). Genome-wide heatmaps and aggregate profiles similarly showed reduced focal CBX2 enrichment and altered RING1B occupancy across their respective called peaks (**Figure 5B**).

**Figure 5.**
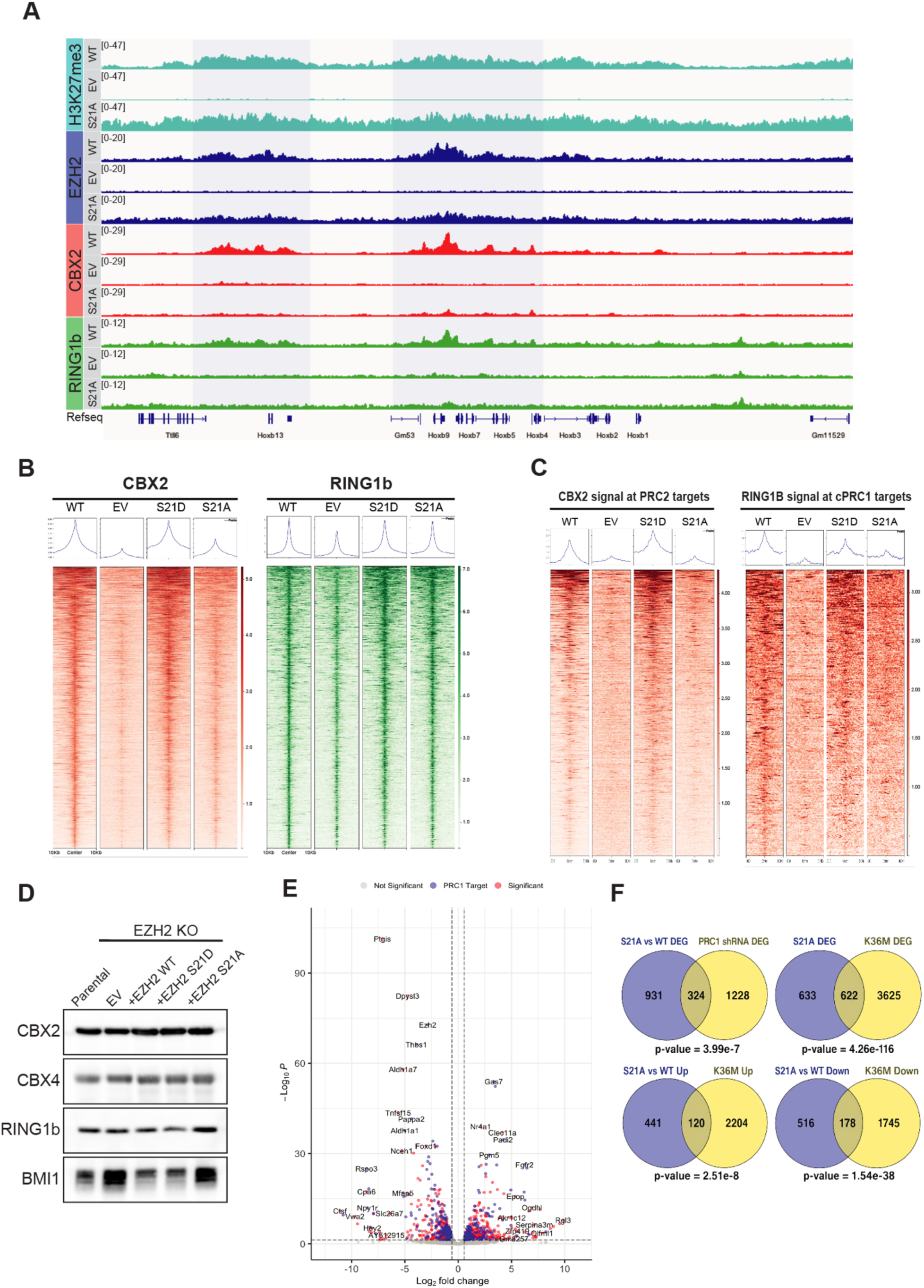
EZH2 S21A reduces focal canonical PRC1 occupancy and alters Polycomb-regulated gene expression. **A.** Genome browser tracks at a representative Polycomb target interval showing ChIP-Rx-normalized H3K27me3, EZH2, CBX2, and RING1B signal in *Ezh2*-knockout C3H10T1/2 cells expressing wild-type EZH2, empty vector, or EZH2 S21A. Shaded regions mark intervals where CBX2 and RING1B signal is reduced or lost in EZH2 S21A-expressing cells relative to wild-type EZH2-expressing cells. **B.** Heatmaps and aggregate profiles of ChIP-Rx-normalized CBX2 and RING1B signal centered on their respective called peaks in *Ezh2*-knockout C3H10T1/2 cells expressing wild-type EZH2, empty vector, EZH2 S21D, or EZH2 S21A. Each row represents an individual peak ordered by signal intensity, and the plots span 10 kb on either side of the peak center. **C.** Heatmaps and aggregate profiles of ChIP-Rx-normalized CBX2 signal at PRC2 target sites and RING1B signal at canonical PRC1 target sites in cells expressing the indicated EZH2 constructs. PRC2 target sites were defined as genomic intervals with overlapping EZH2 and H3K27me3 peaks. Canonical PRC1 target sites were defined as intervals with overlapping EZH2, H3K27me3, and CBX2 peaks. Heatmaps illustrate redistribution and reduced focal CBX2 and RING1B signal in EZH2 S21A-expressing cells. **D.** Immunoblot analysis of parental C3H10T1/2 mesenchymal cells and *Ezh2*-knockout cells reconstituted with empty vector, wild-type EZH2, EZH2 S21D, or EZH2 S21A. Blots were probed for the canonical PRC1 subunits CBX2, CBX4, RING1B, and BMI1. **E.** Volcano plot of differentially expressed genes between EZH2 S21A- and wild-type EZH2-expressing cells. Genes meeting an absolute fold-change threshold greater than 1.5 and q-value less than 0.05 are shown in red. Genes within this set that were also differentially expressed after RING1B knockdown relative to a scrambled siRNA control are shown in blue, indicating overlap between EZH2 S21A-dependent and canonical PRC1-dependent transcriptional programs. **F.** Venn diagrams showing overlap between EZH2 S21A-regulated genes and genes altered after PRC1 shRNA-mediated depletion or in H3K36M-expressing cells. The lower diagrams separately compare upregulated and downregulated gene sets in EZH2 S21A- and H3K36M-expressing cells. P values were calculated by one-sided Fisher exact tests using as background the genes detected and tested in all relevant RNA-seq comparisons after filtering.

To assess canonical PRC1 redistribution more directly, we examined CBX2 and RING1B signal at defined Polycomb target classes. PRC2 target sites were defined by overlapping EZH2 and H3K27me3 peaks, whereas canonical PRC1 target sites were defined by overlapping EZH2, H3K27me3, and CBX2 peaks in wild-type EZH2-expressing cells. At these regions, S21A caused a marked reduction in CBX2 enrichment and depletion of RING1B at canonical PRC1 targets (**Figure 5C**). Immunoblot analysis showed that steady-state levels of canonical PRC1 subunits were not detectably altered (**Figure 5D**), indicating that reduced CBX2 and RING1B occupancy reflects altered chromatin localization rather than loss of PRC1 protein abundance. These data support a model in which S21A-driven H3K27me3 spreading expands the pool of H3K27me3-marked chromatin, reduces CBX2 and RING1B enrichment at high-occupancy Polycomb target loci, and may redistribute canonical PRC1 across a broader H3K27me3 landscape. This conclusion is consistent with prior evidence that the local abundance and spatial distribution of H3K27me3, together with CBX-dependent recognition of this mark, shape canonical PRC1 occupancy and PRC1-mediated long-range interactions among Polycomb target loci^60,64–67^. Thus, both loss of the H3K27me3-dependent recruitment signal, as in *Ezh2*-knockout cells expressing empty vector, and excessive H3K27me3 spreading, as in S21A-expressing cells, can impair focal canonical PRC1 enrichment at Polycomb target loci, but through distinct mechanisms.

### EZH2 S21A links altered Polycomb targeting to defective mesenchymal gene control

Having found that EZH2 S21A broadens H3K27me3 deposition and redistributes canonical PRC1 from Polycomb target loci, we next asked how these chromatin changes affect gene expression programs that support mesenchymal differentiation. RNA-seq analysis identified broad transcriptional differences between wild-type EZH2- and EZH2 S21A-expressing cells (**Figure 5E**). Within the differentially expressed gene set, genes also altered after RING1B knockdown were highlighted, indicating overlap between EZH2 S21A-dependent and canonical PRC1-dependent transcriptional programs (**Figure 5E**). Gene ontology analysis showed enrichment of developmental and morphogenetic pathways among downregulated genes, whereas upregulated genes were enriched for categories associated with epithelial cell proliferation and related regulatory programs (**Figure S5B**). We compared the S21A transcriptional profile with RNA-seq data from C3H10T1/2 cells expressing H3 K36M, which increases H3K27me3 spreading and impairs mesenchymal differentiation^21^. Because both perturbations broaden H3K27me3 deposition, we asked whether they converge on shared transcriptional outputs. Venn analyses showed a statistically significant overlap between EZH2 S21A- and H3 K36M-regulated genes, including a subset altered in the same direction across both contexts (**Figure 5F**). However, many overlapping genes changed in opposite directions, indicating that S21A and H3 K36M converge on increased H3K27me3 spreading without producing equivalent transcriptional states. This distinction is consistent with mechanistically different routes to altered H3K27 methylation and Polycomb regulation.

Because S21A also reduced CBX2 and RING1B occupancy at canonical PRC1 target loci, we compared genes differentially expressed in S21A-expressing cells with genes altered after PRC1 shRNA-mediated depletion. These datasets showed a statistically significant overlap (**Figure 5F**), linking the transcriptional changes observed in S21A-expressing cells to compromised canonical PRC1 function. Together with the chromatin profiling, these analyses indicate that S21A-driven H3K27me3 spreading and canonical PRC1 redistribution are accompanied by broad disruption of mesenchymal transcriptional programs, providing a mechanistic link between altered Polycomb regulation and impaired lineage differentiation.

## Discussion

Here, we identify phosphorylation of EZH2 S21 as a post-translational input that restrains PRC2 allosteric amplification. Across structural, biochemical, and cellular assays, the phospho-null S21A substitution increased basal catalytic activity, reduced the activator-peptide concentration required for H3K27me3-dependent stimulation, and shifted activator-bound PRC2 toward the compact active conformation. These data support a model in which S21-phos does not block EED-mediated allosteric priming, but instead limits the probability that primed PRC2 completes the transition into the compact, allosterically engaged active state. In cells, loss of this phosphorylation input broadened H3K27me3 deposition beyond initial PRC2 recruitment sites, reduced focal canonical PRC1 enrichment at Polycomb target loci, disrupted mesenchymal gene-control programs, and impaired adipogenic and chondrogenic differentiation. Thus, EZH2 S21-phos tunes the strength of PRC2 positive feedback after chromatin recruitment, rather than acting primarily through changes in complex abundance, accessory subunit composition, or bulk chromatin association. This model is summarized at the enzyme level in **Figure 6A** and at Polycomb target loci in **Figure 6B**.

**Figure 6.**
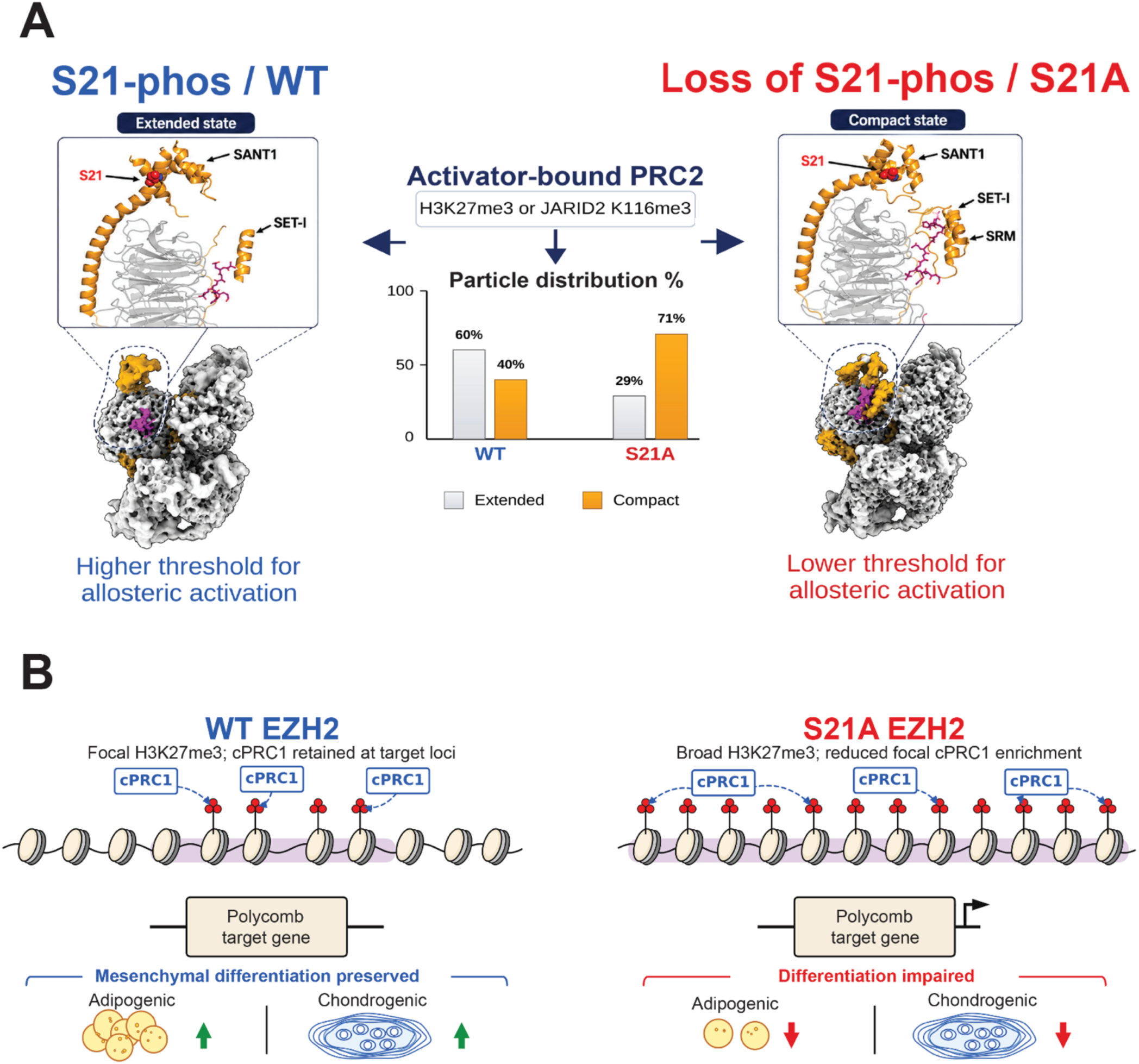
Model for EZH2 S21 phosphorylation-dependent restraint of PRC2 allosteric activation and H3K27me3 propagation. **A.** Model for how EZH2 S21 phosphorylation tunes the allosteric activation threshold of activator-bound PRC2. EED-dependent binding of trimethyl-lysine activator ligands, including product-derived H3K27me3 and cofactor-derived JARID2 K116me3, places PRC2 in an allosterically primed state that can sample extended active and compact active conformations. In the purified WT PRC2 preparation, which contains S21-phosphorylated molecules but is not assumed to be uniformly phosphorylated, cryo-EM particle classification favors the extended active state. This conformational bias is consistent with S21 phosphorylation acting as a restraint that limits conversion of allosteric priming into the compact active, allosterically activated conformation. In phospho-null EZH2 S21A PRC2, loss of the S21 phosphorylation input shifts most particles toward the compact active conformation, consistent with a lower threshold for compact-state activation and enhanced responsiveness to activator peptide stimulation. **B.** WT EZH2-expressing cells are modeled as containing a heterogeneous pool of PRC2 complexes, including molecules with and without AKT1-mediated EZH2 S21 phosphorylation. This mixed phosphorylation state is proposed to tune the effective allosteric set point of PRC2 after recruitment, allowing H3K27me3 propagation while restraining excessive domain expansion. Under these conditions, H3K27me3 remains comparatively focal at Polycomb target loci, CBX2- and RING1B-containing canonical PRC1 is retained at H3K27me3- marked chromatin, and adipogenic and chondrogenic differentiation potential is preserved. In cells expressing phospho-null EZH2 S21A, this phosphorylation-sensitive restraint is unavailable, increasing compact-state sampling and PRC2 responsiveness to local H3K27me3. The resulting excess H3K27me3 deposition broadens H3K27me3 domains from sites of PRC2 recruitment and reduces or redistributes focal canonical PRC1 enrichment at high-occupancy Polycomb target loci. These chromatin changes are associated with misexpression of Polycomb-regulated genes and impaired mesenchymal lineage differentiation.

A central implication of this work is that the relationship between EED ligand binding, allosteric priming, and compact-state activation is not a fixed property of the enzyme complex. H3K27me3 recognition by EED provides a product-stimulated feedback mechanism that can reinforce and spread H3K27 methylation, and methylated cofactors such as JARID2 K116me3 and PALI1 K1241me3 can engage related EED-dependent stimulatory pathways ^12,30,32,33,53^. Recent work further indicates that cofactor-encoded H3K27me3-mimetic inputs can also restrain excessive spreading, emphasizing that the PRC2 read-write loop is actively modulated rather than allowed to propagate without constraint^68^. Our data add phosphorylation of a core PRC2 subunit to this regulatory logic. To our knowledge, EZH2 S21-phos is the first phosphorylation event on a core PRC2 subunit shown to attenuate the dose-response to H3K27me3-derived allosteric activator peptide. Thus, S21 phosphorylation is best viewed not as a simple inhibitor of PRC2, but as a rheostat for allosteric sensitivity: when S21 is phosphorylated, PRC2 is less readily driven into the compact active state; when S21-phos is absent, the same allosteric priming input produces greater catalytic output and enhanced H3K27me3 spreading.

This mechanism provides a route by which PRC2 responsiveness may be coupled to cellular state. EZH2 S21 was originally identified as an AKT phosphorylation site, and AKT is a central effector of PI3K signaling downstream of growth factor, nutrient, metabolic, and stress-responsive inputs ^44,69,70^. The PI3K/AKT pathway is frequently altered in cancer through changes in PIK3CA, PTEN, AKT, mTOR, receptor tyrosine kinase signaling, and related pathway components, suggesting a setting in which S21 phosphorylation could become uncoupled from normal developmental or metabolic regulation ^71^. In this model, increased S21 phosphorylation dampens PRC2 responsiveness to H3K27me3-derived stimulation, whereas reduced S21 phosphorylation lowers the threshold for compact-state activation and H3K27me3 spreading. Signaling pathways may therefore influence not only where PRC2 is recruited but also how strongly it responds to the chromatin marks that activate it.

This model helps explain previous observations that S21 phosphorylation reduces H3K27 methylation, while providing a mechanism for how this inhibition is achieved ^42,44,45,51^. In our assays, S21A increased PRC2 activity without measurably increasing bulk DNA or nucleosome binding and without detectably altering PRC2 complex composition. Instead, S21A increased the fraction of PRC2 in the compact active state and enhanced responsiveness to H3K27me3-derived activator peptide. The gain-of-activity phenotype was suppressed when the EED-EZH2 allosteric interface was disrupted, indicating that S21 phosphorylation regulates PRC2 through the canonical EED-EZH2 allosteric coupling pathway rather than through an independent effect on substrate binding, complex assembly, or catalytic specificity. Thus, S21 phosphorylation acts as a negative gate on PRC2 positive feedback: it does not abolish PRC2 activity but limits how readily local H3K27me3 is converted into additional H3K27 methylation.

The structural and biochemical data support a mechanism in which phosphorylation shifts a pre-existing conformational distribution rather than creating a distinct PRC2 architecture. Wild-type and S21A PRC2 both sample compact and extended active states, and the EZH2 SET domain remains closely similar across these reconstructions. The main S21A-dependent effect is concentrated in the EZH2 regulatory module that connects EED ligand recognition and allosteric priming to compact-state activation. In the S21A complex, this module is biased toward the compact active state, in which SBD bending, SRM ordering, and positioning of SANT1 against the catalytic core are hallmarks of an allosterically activated enzyme. More generally, phosphorylation can perturb local conformational energy landscapes and helix orientation through electrostatic reorganization, raising the possibility that S21-phos biases the SBD toward a straighter extended-active geometry rather than producing a distinct PRC2 architecture^72,73^. This interpretation is consistent with prior structural work linking PRC2 allostery to rearrangements within SANT1, the SBD, the SRM, and the SAL^14,34,35,37^. Importantly, the S21A gain-of-activity phenotype is lost when the EED-EZH2 allosteric interface is disrupted, indicating that S21 phosphorylation regulates PRC2 through the canonical EED-EZH2 allosteric coupling pathway rather than through an independent catalytic mechanism.

The chromatin-level consequences of this altered allosteric set point are not limited to increased H3K27me3 at pre-existing PRC2-bound sites. In both MEFs and mesenchymal progenitors, EZH2 S21A increases H3K27me3 with comparatively modest changes in EZH2 occupancy, supporting a model in which loss of S21-phos increases catalytic output after recruitment rather than primarily increasing recruitment itself (**Figure 6B**). In mesenchymal cells, this expanded H3K27me3 signal is accompanied by reduced CBX2 and RING1B enrichment at canonical PRC1 target loci, despite preserved steady-state levels of PRC1 subunits. These observations suggest that excessive H3K27me3 spreading expands the pool of methylated chromatin and redistributes canonical PRC1 away from high-occupancy Polycomb target regions. This model is consistent with prior work showing that CBX proteins read H3K27me3 in conjunction with additional chromatin features, and that the density and spatial distribution of H3K27me3 influence canonical PRC1 occupancy and long-range interactions among Polycomb-regulated loci^21,60–62,66,74^.

The gene expression and differentiation phenotypes indicate that this phosphorylation-dependent restraint is relevant to developmental gene control. In mesenchymal progenitor cells, S21A increases H3K27 methylation, impairs adipogenic and chondrogenic differentiation, and disrupts lineage-associated transcriptional programs. Importantly, S21A retained more differentiation than empty-vector cells, indicating that these phenotypes are not equivalent. Empty-vector cells fail to restore EZH2-dependent H3K27me3 and therefore cannot support normal focal canonical PRC1 recruitment, whereas S21A cells retain EZH2 activity but broaden H3K27me3 domains and reduce focal canonical PRC1 occupancy. Both failure to restore PRC2 activity and excessive H3K27me3 spreading can therefore impair lineage progression through distinct Polycomb defects. Comparison with H3 K36M-expressing cells indicates that both perturbations broaden H3K27me3 deposition and impair mesenchymal differentiation, yet they do not produce identical gene expression states. This distinction suggests that different routes to enhanced H3K27me3 spreading can converge on defective lineage control while preserving mechanistic specificity. The overlap between S21A-regulated genes and genes altered after RING1B depletion further links the gene expression phenotype to compromised canonical PRC1 function. Thus, excessive PRC2 allosteric amplification can paradoxically weaken Polycomb-mediated gene control by redistributing the reader complex that interprets H3K27me3.

Together, these findings support a model in which EZH2 S21-phos functions as a signaling-sensitive rheostat for PRC2 feedback strength. Rather than acting as an on-off switch for PRC2 recruitment or activity, S21-phos adjusts how readily allosterically primed PRC2 adopts the compact, allosterically activated conformation and how strongly it responds to product-derived stimulation. This mechanism provides a means to restrain H3K27me3 spreading, preserve canonical PRC1 localization at Polycomb target loci, and maintain mesenchymal differentiation competence. More broadly, these results suggest that post-translational modifications on chromatin-modifying enzymes can regulate not only catalytic activity but also the sensitivity of feedback loops that propagate chromatin states across the genome. Future studies should define the pathways that control EZH2 S21-phos across developmental, metabolic, and oncogenic signaling states, determine how S21-phos is integrated with other PRC2 modifications and cofactors, and test whether analogous regulation of allosteric responsiveness operates in other chromatin-modifying complexes.

## RESOURCE AVAILABILITY

### Lead contact

Further information and requests for resources and reagents should be directed to and will be fulfilled by the lead contact, Peter Lewis.

### Materials availability

Plasmids generated in this study will be deposited at Addgene. Cell lines generated in this study are available from the lead contact upon reasonable request and completion of any required institutional material transfer agreements.

### Data and code availability

ChIP-seq data generated in this study have been deposited in the NCBI Gene Expression Omnibus under accession number GSE312458. RNA-seq data generated in this study have been deposited in the NCBI Gene Expression Omnibus under accession number GSE312314. No GEO SuperSeries accession has been created. No new atomic coordinate models were deposited; previously determined PDB 6C23, PDB 6C24, and PDB 6WKR models were rigid-body fit into the experimental cryo-EM maps for figure presentation. This study did not generate custom code.

## EXPERIMENTAL MODEL AND STUDY PARTICIPANT DETAILS

### Mammalian cell lines

HEK293T and human 293T cells used for lentivirus production and transient expression assays were obtained from ATCC (CRL-3216; RRID:CVCL_0063). Immortalized C3H10T1/2 murine mesenchymal cells were obtained from ATCC (CCL-226; RRID:CVCL_0190). *Ezh2*-knockout C3H10T1/2 cells and *Ezh2*-knockout MEFs were generated in this study by CRISPR/Cas9 editing and validated by immunoblotting and genomic DNA Sanger sequencing as described below. *Eed*-knockout MEFs were generated from *Eed* ^fl/fl^ MEFs with loxP sites flanking *Eed* exons 3-6. All mammalian cell lines were monitored monthly for mycoplasma contamination using the e-Myco plus Mycoplasma PCR Detection Kit.

### Insect cells and bacterial strains

Sf9 insect cells were obtained from ATCC (CRL-1711; RRID:CVCL 0549) and were used for baculovirus-mediated expression of recombinant PRC2 complexes. High Five cells (BTI-Tn-5B1-4) were obtained from Thermo Fisher Scientific/Invitrogen (B85502; RRID:CVCL C190; ATCC CRL-10859) and were used for PRC2.2 expression. *E. coli* Rosetta(DE3) cells were obtained from MilliporeSigma/Novagen (Cat# 70954) and used for recombinant histone expression.

## METHOD DETAILS

### Cell culture and mycoplasma monitoring

HEK293T, human 293T, and immortalized C3H10T1/2 cells were maintained in high-glucose Dulbecco’s Modified Eagle Medium (DMEM; Sigma-Aldrich, D6429) supplemented with 10% fetal bovine serum (Omega Scientific, FB-11), 100 U/mL penicillin, and 100 μg/mL streptomycin (Life Technologies/Gibco, 15140-163). Cells were maintained on tissue-culture-treated culture dishes, passaged every 2-3 days using 0.25% trypsin-EDTA (Thermo Fisher Scientific/Gibco, 25200072), and cultured at 37°C in a 5% CO2 incubator. Cell lines were monitored monthly for mycoplasma contamination using the e-Myco plus Mycoplasma PCR Detection Kit (Bulldog Bio/LiliF Diagnostics, 25234).

### Generation of transgenic mammalian cell lines

Lentiviruses were generated by co-transfecting HEK293T cells with psPAX2 (Addgene plasmid #12260; RRID:Addgene 12260), pMD2.G (Addgene plasmid #12259; RRID:Addgene 12259), and pCDH-EF1α-MCS-PuroR transfer plasmids using PEI MAX DNA transfection reagent (Polysciences/Kyfora Bio, 24765). The pCDH-EF1α-MCS-PuroR backbone corresponds to the pCDH-EF1α-MCS-IRES-Puro lentivector family (System Biosciences, CD532A-2; Fisher NC2011214), modified as needed to express EZH2 transgenes.

Virus-containing medium was collected 48 h and 72 h after transfection. Human 293T, C3H10T1/2, or MEF cells were transduced with recombinant lentivirus and selected with 1 μg/mL puromycin (GoldBio, P-600-1) for 4 days. Transduced cells were harvested for immunoblot analysis 6-10 days after transduction unless otherwise indicated.

### Generation of CRISPR-edited mammalian cell lines

CRISPR editing plasmids were introduced using PEI MAX DNA transfection reagent (Polysciences/Kyfora Bio, 24765). Cells were transfected with pSpCas9(BB)-2A-Puro (PX459; Addgene plasmid #48139; RRID:Addgene 48139) carrying the appropriate sgRNA sequence. After 24 h, cells were selected with 1 μg/mL puromycin for 4 days. Single cells were isolated into 96-well plates. Successful CRISPR knockouts were screened by immunoblotting with the indicated antibodies and by Sanger sequencing of genomic DNA.

### Mesenchymal differentiation

For adipogenic differentiation, confluent C3H10T1/2 cells were stimulated in DMEM containing 10% FBS, 0.5 mM isobutylmethylxanthine (Sigma-Aldrich, I5879), 1 μM dexamethasone (Sigma-Aldrich, D4902), 5 μg/mL insulin (Sigma-Aldrich, 91077C-100MG), and 5 μM troglitazone (Sigma-Aldrich, T2573). After 2 days of differentiation, cells were maintained in DMEM containing insulin until analysis. For chondrogenic differentiation, confluent cells were stimulated in DMEM containing 1% FBS, 10 μg/mL insulin (Sigma-Aldrich, 91077C-100MG), 3 × 10^−8^ M sodium selenite (Sigma-Aldrich, S5261), 10 μg/mL human transferrin (Sigma-Aldrich, T8158-100MG), 10^−7^ M dexamethasone (Sigma-Aldrich, D4902), and 100 ng/mL rhBMP-2 (R&D Systems/Bio-Techne, 355-BM; distributed by VWR, 103012-266). Medium was replaced with fresh differentiation cocktail every 2-3 days until analysis.

### Alcian blue and Oil Red O staining

For Alcian blue staining, cells were washed in PBS and fixed for 5 min at room temperature with 4% formaldehyde. Cells were stained with 1% Alcian blue 8GX (Sigma-Aldrich, A5268), pH 1.0, for 1 h at room temperature. For quantification, cells were lysed in 1% SDS, and absorbance was measured at 605 nm. For Oil Red O staining, cells were washed in PBS and fixed for 20 min at room temperature with 3% formaldehyde. Cells were stained with Oil Red O (Sigma-Aldrich, O0625). For quantification, Oil Red O staining was dissolved in isopropanol, and absorbance was measured at 500 nm.

### FLAG affinity purification from mammalian nuclear extracts

Nuclei were isolated by resuspending cells in buffer A (15 mM HEPES, pH 7.9, 4 mM MgCl2, 10 mM KCl, 1 mM EDTA, and 0.4 mM PMSF). Nuclei were resuspended in buffer AC (15 mM HEPES, pH 7.9, 110 mM KCl, 4 mM MgCl2, 1 mM EDTA, 0.4 mM PMSF, and 1x protease inhibitor cocktail). Nuclear extract was prepared by adding 0.1 volume saturated ammonium sulfate followed by ultracentrifugation at 85,000 x g for 90 min.

Supernatant was dialyzed against FLAG-IP buffer (20 mM HEPES, pH 7.9, 250 mM KCl, 1 mM EDTA, 2 mM β-mercaptoethanol, 0.4 mM PMSF, and 0.1% Triton X-100). Nuclear extract was incubated with M2 anti-FLAG affinity gel (Sigma-Aldrich, A2220-2X25ML) for 2 h. Beads were washed three times with wash buffer (20 mM HEPES, pH 7.9, 500 mM KCl, 1 mM EDTA, 2 mM β-mercaptoethanol, 0.4 mM PMSF, and 0.1% Triton X-100). Captured proteins were eluted in 20 mM HEPES, pH 7.9, 200 mM KCl, 1 mM EDTA, 10% glycerol, 0.4 mM PMSF, and 500 μg/mL 3×FLAG peptide (Tufts University Core Facility, custom synthesis).

### Immunoblotting

Immunoprecipitation samples or whole-cell lysates were separated by SDS-PAGE, transferred to nitrocellulose membranes, blocked in 5% nonfat milk in Tris-buffered saline containing 0.5% Tween-20, and probed with primary antibodies at 1:1,000 unless otherwise indicated. Horseradish peroxidase-conjugated anti-rabbit secondary antibody (Bio-Rad, 1706515) or anti-mouse secondary antibody (Jackson ImmunoResearch, 115-035-003) was used at 1:5,000. Antibodies are listed in the Key Resources Table.

### Histone extraction

Cells were lysed in hypotonic lysis buffer containing 10 mM HEPES, 10 mM KCl, 1.5 mM MgCl_2_, 0.5 mM PMSF, and 0.1% Triton X-100. Chromatin pellets were incubated overnight at room temperature in 0.4 N H_2_SO_4_. Histones were precipitated from the supernatant using 33% trichloroacetic acid, washed in acetone, and resuspended in water.

### Histone derivatization and PTM analysis by nano-LC-MS

Acid-extracted histones (15 μg) were resuspended in 100 mM ammonium bicarbonate (pH 8) and derivatized using propionic anhydride similar to previous studies^75^. Two volumes of histone were added to one volume of 25% propionic anhydride diluted in 2-propanol. Extra ammonium bicarbonate salt was added to buffer the pH. After incubation at 37°C for 15 mins, the samples were dried and resuspended in 20 μl of 100 mM ammonium bicarbonate for a second round of derivatization. Derivatized histones were resuspended in 20 μl of 100 mM ammonium bicarbonate for digestion with trypsin (1 μg) overnight at room temperature. The propionic anhydride derivatization was then repeated twice to propionylate the newly generated peptide N-termini. Histone peptides were subsequently desalted with C18 stage tips and analyzed by LC-MS/MS. Histone peptides were separated with a Dionex UltiMate 3000 system (Thermo) over a fused silica column (Polymicro Tech, 75 μm i.d. x 12 cm) packed with C18 material (ReproSil-Pur 120 C18-AQ, 3 μm, Dr. Maisch GmbH) and eluting into a QE-HF mass spectrometer (Thermo). Water containing 0.1% formic acid and 80% acetonitrile/20% water containing 0.1% formic acid were used as solvents A and B, respectively. The chromatography gradient consisted of 0 to 5% solvent B over 0 to 4 mins, 5 to 33% solvent B over 4 to 49 mins, 33 to 98% solvent B over 49 to 54 mins, and holding at 98% solvent B for 54 mins to 63.9 mins. The flow rate was set to 300 nl/min. Histone peptides were analyzed by data-independent acquisition (DIA) MS using a cycle of two full MS scans separated by eight DIA MS/MS scans. The cycle consisted of one full MS scan in positive centroid mode with resolution 60,000, scan range 300-1100 m/z, automatic gain control 1e6, and max injection time 50 ms. Next, eight DIA MS/MS scans were acquired with resolution 30,000, automatic gain control 5e5, and auto max injection time. The eight DIA windows used a width of 50 m/z and were centered on 325, 375, 425, 475, 525, 575, 625, and 675 m/z. HCD was used as the fragmentation method with 27 NCE. A second full MS scan was then performed after the first eight DIA scans, which was then followed by an additional eight DIA MS/MS scans centered on 725, 775, 825, 875, 925, 975, 1025, and 1075 m/z. The relative abundances of histone modifications were calculated based on chromatographic peak area using EpiProfile.

### Native oligonucleosome purification

Native oligonucleosomes were purified from *Eed* KO Mouse Embryonic Fibroblasts. Nuclei were prepared by resuspending 100 million cells in buffer-A and centrifugation at 3000 RPM for 5 min. Nuclei were resuspended in buffer-AP (15 mM HEPES pH 8, 15 mM NaCl, 60mM KCl, 5% Sucrose, 0.5 mM spermine, 0.15 mM spermidine, 0.4 mM PMSF, 1mM β-mercaptoethanol) and treated with 0.2 units/µl MNase for 20 min at 37° C. After quenching with 5 mM EDTA, nuclei were centrifuged at 3000 RPM for 5 min. Nuclei were lysed by resuspension in 10 mM EDTA and 500 mM NaCl. Oligonucleosomes were purified over sucrose gradient (5-30% sucrose, 15mM HEPES pH7.9, 1 mM EDTA, 500 mM NaCl, 5 mM PMSF). Oligonucleosomes in fractions 15-21 ml were concentrated and dialyzed against 15 mM Tris pH 8.0, 100 mM NaCl, 1 mM EDTA, 4 mM PMSF, 10% glycerol.

### Purification of recombinant core PRC2

Recombinant PRC2 complex was purified from SF9 cells co-infected with baculoviruses containing human FLAG-tagged Ezh2, Suz12, EED and Rbbp4. Cells were lysed in lysis buffer (15 mM Tris pH8.0, 5 mM MgCl2, 500 mM KCl, 0.5% Triton X-100, 1 mM EDTA, 0.4 mM PMSF, 1X Protease Inhibitor Cocktail, β-mercaptoethanol), followed by M2-FLAG affinity purification and anion-exchange chromatography.

### Purification of PRC2.2 WT and S21A (AJ106-350)

A multibac plasmid containing sequences for human EZH2, SUZ12 (aa 70-685), EED (aa 80-441), RBAP48/RBBP4, strepIIGFP-AEBP2 embryonic isoform, and strepIIGFP-JARID2 (106-350) was assembled using LIC cloning (Berkeley MacroLab). Once the multibac plasmid was assembled, the S21A EZH2 point mutation was generated by Genscript. Sf9 or HighFive cells were infected with baculovirus and expressed for 66 hours. Cells were lysed in lysis buffer (25mM HEPES pH=7.9, 250mM NaCl, 1mM MgCl2, 0.1% NP-40, 0.2 mM PMSF, and 1X Protease Inhibitor Cocktail, 1mM TCEP), affinity purified with Streptactin XT resin (IBA), TEV cleaved overnight and further purified with heparin chromatography and size-exclusion chromatography (Superose 6 10/300).

### Cryo-EM sample preparation

Homogeneous wild-type and EZH2 S21A PRC2 complexes were assessed by negative-stain EM for particle integrity. For cryo-EM preparation, S-adenosyl-L-homocysteine (Sigma-Aldrich, A9384) was added to PRC2 complexes at 50 μM, and complexes were crosslinked with 0.5 mM BS3 (Thermo Fisher Scientific/Pierce, 21580) for 30 min. Crosslinked complexes were desalted using a 75-μL Zeba Micro Spin Desalting Column (Thermo Fisher Scientific/Pierce, 89877) into 25 mM HEPES, pH 7.9, 75 mM NaCl, 1 mM TCEP, and 5% glycerol. A final cryo-EM buffer containing 2% trehalose and 0.01% β-octyl glucoside was used. Crosslinked PRC2 was diluted to 400 nM in cryo-EM buffer and applied immediately to glow-discharged Quantifoil R 2/1 300-mesh gold grids (Quantifoil/Electron Microscopy Sciences, Q350AR1; Fisher NC1523189) for 30 s. Samples were vitrified in liquid ethane using a Vitrobot Mark IV after a 3.5-s blot.

### Cryo-EM data collection and processing

Roughly 15k to 16k micrographs of the S21A and wild-type PRC2 each were collected on a Titan Krios G3i 300 keV equipped with a Falcon4i Ceta-D camera and a Selectris X energy filter. The physical pixel size was 0.7551 Å/pix and data were collected in EER format movies. Patch motion correction and patch CTF were performed in CryoSPARC. The general processing scheme was based on ‘Case Study: End-to-end and exploratory processing of a motor-bound nucleosome (EMPIAR-10739)’ found within the CryoSPARC guide. Bad micrographs were discarded using the micrograph junk detector and manually curated exposure tools. Micrographs were subsequently denoised, and the blob picker tool was applied to pick an initial set of particles for template generation. Following multiple rounds of 2D classification, a set of 2D classes of PRC2 was used for template picking. Once picks were inspected, particles were extracted at a box size of 384 px and downsampled to 64px. An initial heterorefinement of particles picked was performed to remove junk. One class yielded an intact PRC2 particle (∼30% of particles), and that class of particles was then re-extracted at 384px box size. Using the subset particles by statistics tool in CryoSPARC, we identified a smaller population among the extracted particles that yielded high-resolution refinements of PRC2. The final sorted 199k or 300k particles, for each of the data sets respectively, were subjected to heterogeneous refinement with two input volumes of EMD-7334 and EMD-7335, which were low-pass filtered to 20 Å and 30 Å. Following classification, all particles were then subjected to a non-uniform refinement (Punjani et al. 2020) and reference-free 2D class averages. All reported resolutions are based on the gold-standard FSC=0.143 criterion^76^. Quality of each refinement and maps was assessed by GSFSC within CryoSPARC and then subsequently with the 3DFSC server (New York Structural Biology Center and Salk Institute). Sphericity and global resolution of each refined map are reported in the supplemental figure 3. The final wild-type and S21A refinement maps and half-maps were deposited to the EMDB (EMD-76889, EMD-76893, EMD-76902, EMD-76903) along with their locally filtered maps (B-factor applied = -10). Rigid-body docking of PDB-6C23 and PDB-6C24 into these maps was used for the figure presentation. All figures were generated in Chimera and ChimeraX.

### Reconstitution of recombinant nucleosomes

Recombinant histones were purified from E. coli Rosetta cells. Briefly, inclusion bodies were solubilized using 6.3 M Guanidine-HCl, 500 mM NaCl, 50 mM Tris pH 8 and 10 mM β-mercaptoethanol. Histones were purified using Ni-NTA chromatography. Guanidine-HCl was removed using PD10 desalting column and histones were lyophilized overnight. Lyophilized histones were resuspended in D0 buffer (50 mM Tris pH 8.5, 6.3 M Guanidine-HCl, 4 mM EDTA, 10 mM β-mercaptoethanol) and dialyzed against Refolding buffer (20 mM Tris pH 7.5, 1 mM EDTA, 2 M NaCl, 5 mM DTT) for 48 hours with two changes of buffer. Octamers were purified using Superdex 200 column. To reconstitute nucleosomes, octamers and DNA were mixed at 1:1 ratio and dialyzed against 20 mM Tris pH 7.5, 1.4 M NaCl, 1 mM EDTA, 10 mM β-mercaptoethanol for 2 hours. Salt was gradually reduced to 100 mM overnight by pumping Ending buffer at 1ml/min (20 mM Tris pH 7.5, 10 mM NaCl, 1 mM EDTA, 10 mM β-mercaptoethanol). Nucleosomes were finally dialyzed against Ending buffer and stored at -80oC in 10% glycerol. Cy5 and Cy3 labelled DNA 601 for labelled nucleosomes were generated through PCR using labelled forward primers ordered from Sigma.

### Histone Methyltransferase assays

For a typical PRC2 reaction, 200 nM oligonucleosome or 25 μM H3 (18-37) peptide substrate was incubated with 20 nM PRC2 complex, 4 μM S-adenosyl Methionine (1 μM ^3H^-SAM (Perkin Elmer); 3 μM cold SAM) and 40 μM H3K27me3 peptide in 2X reaction buffer (50 mM Tris pH 8.0, 4 mM MgSO4, 10 mM DTT and 8 mM PMSF) for 60 min. For scintillation counting, reaction was spotted on a phosphocellulose filter (p81) and dried for 10 min. Filters were washed 3x with 0.1 M NaHCO3 for 30 min, rinsed with acetone and dried for 10 min. Scintillation counting was performed using Tri-Carb 2910 TR liquid scintillation analyzer (Perkin Elmer). Counts were corrected for background using reaction without substrate. For reactions evaluated by western blot analysis, similar conditions were used with ^3^H-SAM excluded from the reaction mixture.

### Electrophoretic mobility shift assays

Binding of recombinant wild-type or EZH2 S21A PRC2 to Cy5-labeled 601 DNA and Cy5-labeled 601 nucleosomes was assessed by EMSA. PRC2 was titrated in two-fold dilutions beginning at 500 nM for DNA-binding assays and 1000 nM for nucleosome-binding assays, with a no-PRC2 control. Fraction bound was quantified from two independent replicates, and apparent Kd values were obtained by nonlinear fitting with Hill slope.

### Chromatin Immunoprecipitation

Cells were cross-linked using 1% paraformaldehyde for 5 min at room temperature and quenched with 200 mM glycine. Approximately 20 million cells were lysed in digestion buffer containing 20 mM Tris, pH 7.6, 1 mM CaCl2, 0.25% Triton X-100, 5 mM PMSF, and 1x protease inhibitor cocktail. Chromatin was treated with 10 units MNase per million cells for 10 min at 37°C and quenched with 5 mM EDTA. Chromatin was solubilized by sonication using a Covaris instrument at 120 peak incident power, 5% duty factor, and 200 cycles/burst for 3 min and dialyzed against RIPA buffer for 2 h. Chromatin concentration was measured using Qubit, and 293T or *Drosophila* S2 spike-in chromatin was added to MEF samples at a 1:40 ratio. Chromatin was incubated overnight with 2-5 μg primary antibody, captured with Dynabeads for 4 h, washed three times with RIPA, three times with RIPA plus 300 mM NaCl, and twice with RIPA-LiCl for 5 min each, and eluted in 10 mM Tris, 1 mM EDTA, and 1% SDS. Eluted DNA was reverse-crosslinked overnight at 65°C, treated with proteinase K, and purified using PCR purification columns. Illumina sequencing libraries were generated using the NEBNext Ultra DNA Library Prep Kit. ChIP experiments were performed in at least two independent replicates with similar results; at least one replicate was sequenced for ChIP-seq, and ChIP-qPCR data are displayed as technical replicate measurements where indicated.

### ChIP-Sequencing analysis

Reads that passed quality score were aligned to mouse (mm9), human (hg38) or Drosophila (dm3) genomes using Bowtie2 (parameters: -q -v 2 -m 2 -a –best –strata) ^77^. Sample normalization factor was determined as ChIP-Rx = 10^6^ / (total reads aligned to exogenous reference genome) or RPKM = 10^6^ / (total aligned reads). SAM files were converted to BAM files using samtools^78^. BigWig files were generated using deeptools (-bs 50 –smoothLength 600)^79^. Peaks were called using MACS2. Statistical analysis was performed using R. Deeptools and IGV genome browser was used for data visualization^79,80^. PRC2 target sites were defined as regions with overlapping EZH2 and H3K27me3 peaks, and canonical PRC1 target sites were defined as regions with overlapping EZH2, H3K27me3, and CBX2 peaks using bedtools v2.30.0. H3K27me3 spreading analyses compared signal at EZH2 recruitment sites with signal in 50-kb flanking regions.

### RT-qPCR

Cells were collected for RNA extraction by centrifugation at 300 x g for 3 min, lysed in TRIzol, and frozen until purification. RNA was purified using the Quick-RNA MiniPrep Kit (Zymo Research, R1055) with off-column DNase I digestion. Purified RNA was stored at -80°C. Reverse transcription was performed with GoScript Reverse Transcription Mix, Random Primers (Promega, A2801). qPCR was performed with the indicated primers and Power SYBR Green PCR Master Mix (Life Technologies/Invitrogen, 4368708) using the following program: 95°C for 10 s, followed by 52 cycles of 95°C for 20 s, 60°C for 30 s, and 72°C for 20 s.

### RNA-sequencing

For each RNA-seq library, 1000 ng RNA was used for library preparation with the NEBNext Poly(A) mRNA Magnetic Isolation Module (New England Biolabs, E7490L), NEBNext Ultra II DNA Library Prep Kit for Illumina (New England Biolabs, E7645L), and NEBNext Multiplex Unique Dual Index Oligos for Illumina (New England Biolabs, E6440S). Libraries were sequenced on an Illumina NovaSeq 6000 using 50-bp paired-end reads to a target depth of 25 million reads per sample. Two biological replicates were generated for each experimental condition.

### RNA-sequencing analysis

Reads that passed quality score were aligned to mouse (mm9) transcriptome using STAR -2.7.8a and the follow parameters (--quantMode TranscriptomeSAM GeneCounts)^81^. For host genes Salmon and DEseq2 were used for differential expression^82^. All analysis were performed with two biological replicates. Data were visualized using ggplot2. Genes with fewer than 10 total counts across all samples were removed before differential-expression testing. DESeq2 Wald tests were used for pairwise comparisons, and p values were adjusted using the Benjamini-Hochberg method. Data were visualized using ggplot2 v3.5.1 in R v4.4.1.

Differentially expressed genes were defined using the thresholds indicated in the figure legends, including fold change greater than 1.5 and q value less than 0.05 for Figure 5E. Gene overlap p values were calculated using one-sided Fisher exact tests with as background the set of genes detected and tested for differential expression in all relevant RNA-seq comparisons after filtering. Gene ontology enrichment analysis was performed on downregulated and upregulated gene sets using clusterProfiler v4.12.6 and org.Mm.eg.db v3.19.1 with the biological process ontology; enrichment p values were adjusted using the Benjamini-Hochberg method.

## QUANTIFICATION AND STATISTICAL ANALYSIS

Statistical details for each experiment, including replicate number, error bars, and statistical tests, are provided in the figure legends where available. Data are presented as mean +/- SEM or mean +/- SD as indicated in the legends. Bar-graph comparisons among three or more groups, including Figures 1D, 4C, 4D, S5C, and related panels, were analyzed by ordinary one-way ANOVA followed by Tukey’s multiple-comparisons test in GraphPad Prism v10.2.3. Two-group bar-graph comparisons were analyzed by two-tailed unpaired t tests unless a paired design or non-parametric test is specified. Nonlinear regression was used to fit H3K27me3 dose-response data, Michaelis-Menten kinetic data, and EMSA binding data. Differential expression analyses were performed using DESeq2 v1.44.0 with Wald tests and Benjamini-Hochberg correction. Gene-overlap analyses were evaluated with one-sided Fisher exact tests using the background gene sets described above. ChIP-qPCR comparisons were evaluated using paired non-parametric tests where specified in the prior laboratory methods. Boxplot comparisons used Wilcoxon rank-sum tests where indicated. p-value symbols are defined as follows: ns, not significant; *p < 0.05; **p < 0.01; ***p < 0.001; ****p < 0.0001.

### Oligonucleotide primers

Primers used for RT-qPCR and ChIP-qPCR are listed below.

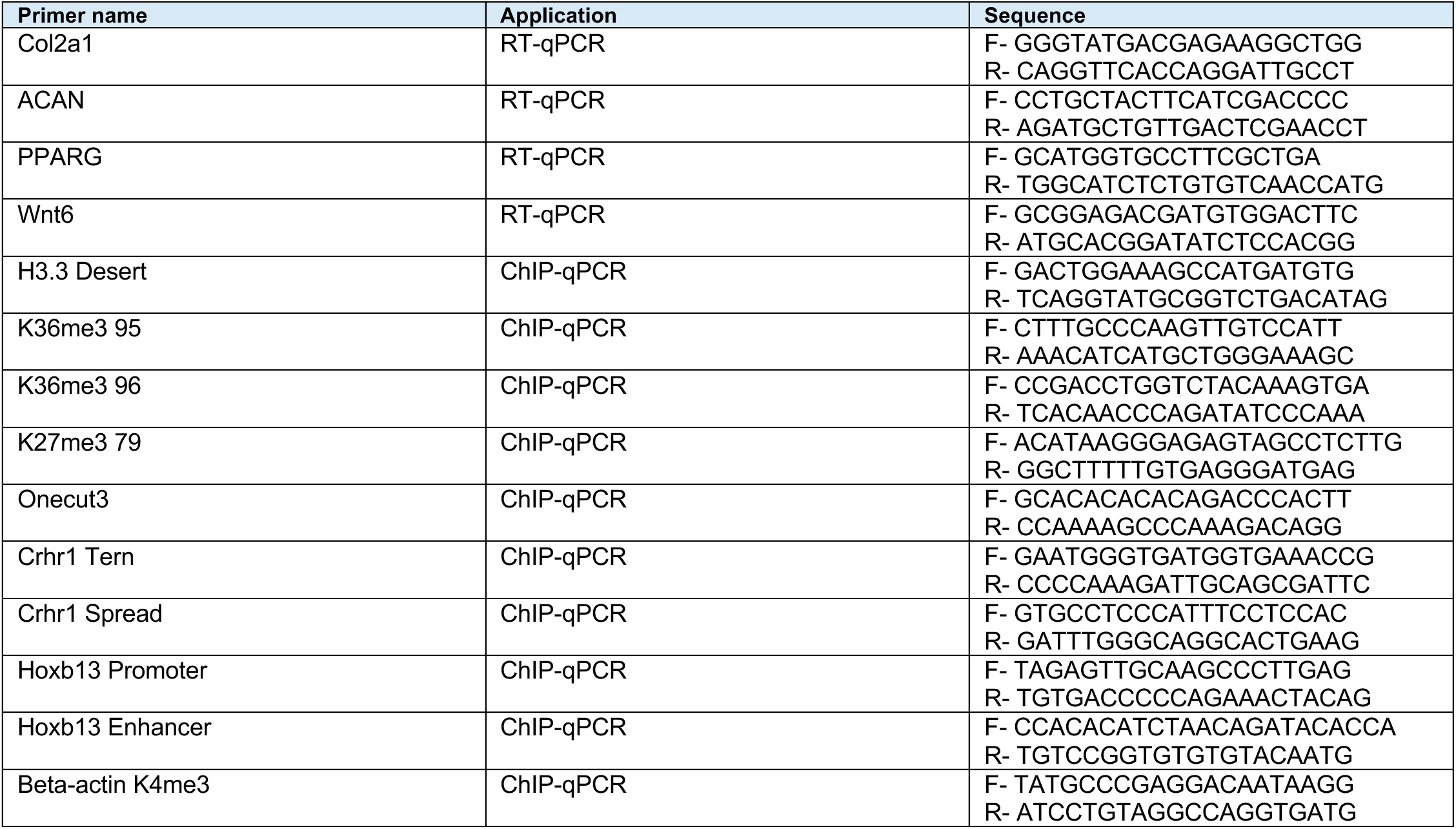

## Acknowledgements

This work was supported by the National Institutes of Health (R01 CA266861 to P.W.L.; P01 CA196539 to P.W.L. and B.A.G.; P30 CA014520 to the UW Carbone Cancer Center; R35 GM155426 to V.K.; and R24 OD033699 to the CU Boulder Biochemistry Shared Instrumentation Pool), the National Science Foundation (MCB 2446197 to V.K.), CU Boulder startup funds to V.K., an Innovator Award from Alex’s Lemonade Stand Foundation to P.W.L., and grants from the Childhood Brain Tumor Foundation and Rally Foundation for Childhood Cancer Research to P.W.L. V.K. and P.W.L. are Pew Scholars in the Biomedical Sciences, supported by the Pew Charitable Trusts. M.N.M. was supported by funds from NIH R35 GM155426 and NSF MCB 2446197. We thank Ian Fries and the Stanford-SLAC Cryo-EM Center (S2C2) for cryo-EM data collection. We also thank Dr. Erik Hartwick and the BioKEM facility at CU Boulder for cryo-EM grid preparation, and the CU Boulder Biochemistry Shared Instrumentation Pool for experimental support. This work utilized the Blanca condo computing resource at the University of Colorado Boulder, which is jointly funded by computing users and the University of Colorado Boulder.

## Author Contributions

A.Q.R., M.M., V.K., and P.W.L. conceived the study. A.Q.R. performed biochemical and cellular experiments, including recombinant PRC2 purification, methyltransferase assays, EMSAs, generation and characterization of EZH2-reconstituted cell lines, immunoblotting, coimmunoprecipitation, ChIP-seq, ChIP-qPCR, RNA-seq, RT-qPCR, mesenchymal differentiation assays, and analysis and visualization of data. M.M. purified PRC2 complexes for cryo-EM studies and performed cryo-EM data processing, classification, map reconstruction, structural analysis, and structural visualization. E.G.P. performed and analyzed quantitative histone post-translational modification mass spectrometry. A.Q.R. and R.H. generated recombinant nucleosomes. B.A.G. supervised the histone post-translational modification mass spectrometry studies and contributed to data interpretation. V.K. supervised cryo-EM, cryo-EM analysis, and structural interpretation. P.W.L. supervised the biochemical, genomic, and cellular studies and directed the overall project. A.Q.R., M.M., V.K., and P.W.L. wrote the manuscript. All authors reviewed and edited the manuscript.

