## Supplemental Figures for "EZH2 Serine 21 Phosphorylation Restrains Compact-State PRC2 Activation and H3K27me3 Propagation"

**Figure S1**

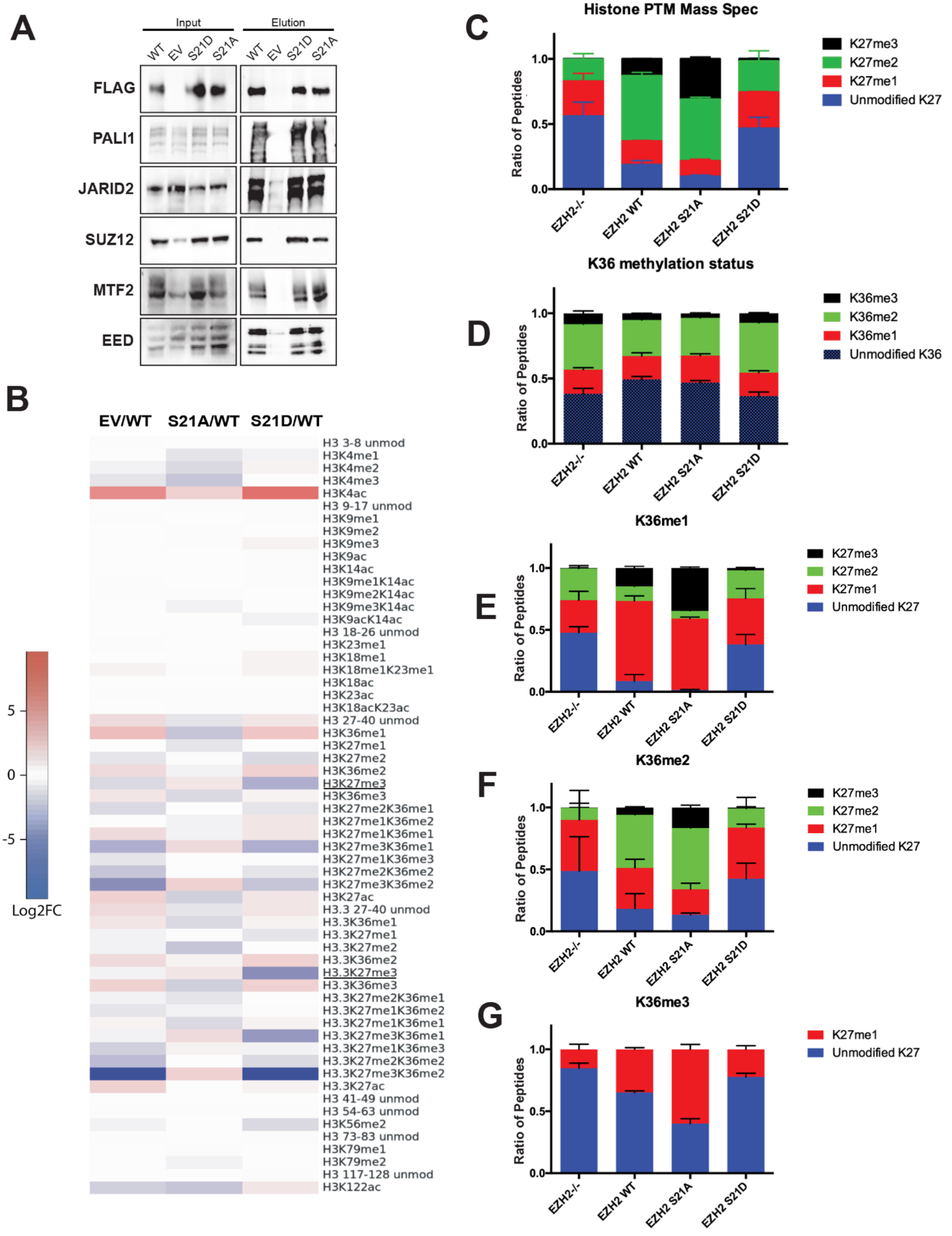
A
Input
Elution
WT
EV
S21D
S21A
FLAG
PALI1
JARID2
SUZ12
MTF2
EED
B
EV/WT
S21A/WT
S21D/WT
H3 3-8 unmod
H3K4me1
H3K4me2
H3K4me3
H3K4ac
H3 9-17 unmod
H3K9me1
H3K9me2
H3K9me3
H3K9ac
H3K14ac
H3K9me1K14ac
H3K9me2K14ac
H3K9me3K14ac
H3K9acK14ac
H3 18-26 unmod
H3K23me1
H3K18me1
H3K18me1K23me1
H3K18ac
H3K23ac
H3K18acK23ac
H3 27-40 unmod
H3K36me1
H3K27me1
H3K27me2
H3K36me2
H3K27me3
H3K36me3
H3K27me2K36me1
H3K27me1K36me2
H3K27me1K36me1
H3K27me3K36me1
H3K27me1K36me3
H3K27me2K36me2
H3K27me3K36me2
H3K27ac
H3.3 27-40 unmod
H3.3K36me1
H3.3K27me1
H3.3K27me2
H3.3K36me2
H3.3K27me3
H3.3K36me3
H3.3K27me2K36me1
H3.3K27me1K36me2
H3.3K27me1K36me1
H3.3K27me3K36me1
H3.3K27me1K36me3
H3.3K27me2K36me2
H3.3K27me3K36me2
H3.3K27ac
H3 41-49 unmod
H3 54-63 unmod
H3K56me2
H3 73-83 unmod
H3K79me1
H3K79me2
H3 117-128 unmod
H3K122ac
5
0
-5
Log2FC
C
Histone PTM Mass Spec
Ratio of Peptides
1.0
0.5
0.0
EZH2-/-
EZH2 WT
EZH2 S21A
EZH2 S21D
K27me3
K27me2
K27me1
Unmodified K27
D
K36 methylation status
K36me3
K36me2
K36me1
Unmodified K36
E
K36me1
F
K36me2
G
K36me3

**Figure S1. Histone H3 post-translational modification profiling and PRC2 complex composition in *Ezh2*-knockout MEFs expressing EZH2 variants, related to Figure 1.**

**A.** Immunoblot analysis of FLAG coimmunoprecipitation from nuclear extracts of *Ezh2*-knockout MEFs expressing FLAG-tagged wild-type EZH2, S21D, S21A, or empty vector. Input and FLAG elution fractions were probed for FLAG, the PRC2 core subunits SUZ12 and EED, the PRC2.1 accessory subunits PALI1 and MTF2, and the PRC2.2 accessory subunit JARID2.

**B.** Heatmap showing $\log_2$ fold changes in the abundance of individual histone H3 post-translational modification states in *Ezh2*-knockout MEFs expressing empty vector, S21A, or S21D relative to cells reconstituted with wild-type EZH2. Rows indicate H3 peptide modification states, and columns indicate the pairwise comparisons EV/WT, S21A/WT, and S21D/WT.

**C.** Quantitative mass spectrometry analysis of H3K27 methylation states in *Ezh2*-knockout MEFs expressing empty vector, wild-type EZH2, S21A, or S21D. Stacked bars show the fractional abundance of H3 peptides containing unmodified K27, H3K27me1, H3K27me2, or H3K27me3. Data are shown as mean ± SEM.

**D.** Quantitative mass spectrometry analysis of H3K36 methylation states in *Ezh2*-knockout MEFs expressing empty vector, wild-type EZH2, S21A, or S21D. Stacked bars show the fractional abundance of H3 peptides containing unmodified K36, H3K36me1, H3K36me2, or H3K36me3. Data are shown as mean ± SEM.

**E-G.** Quantitative analysis of H3K27 methylation states on H3 peptides containing H3K36me1 (**E**), H3K36me2 (**F**), or H3K36me3 (**G**). Stacked bars show the fractional abundance of the indicated K27 methylation states within each H3K36 methylation-defined peptide class in *Ezh2*-knockout MEFs expressing the indicated EZH2 transgenes or empty vector. Data are shown as mean ± SEM.

### Figure S2

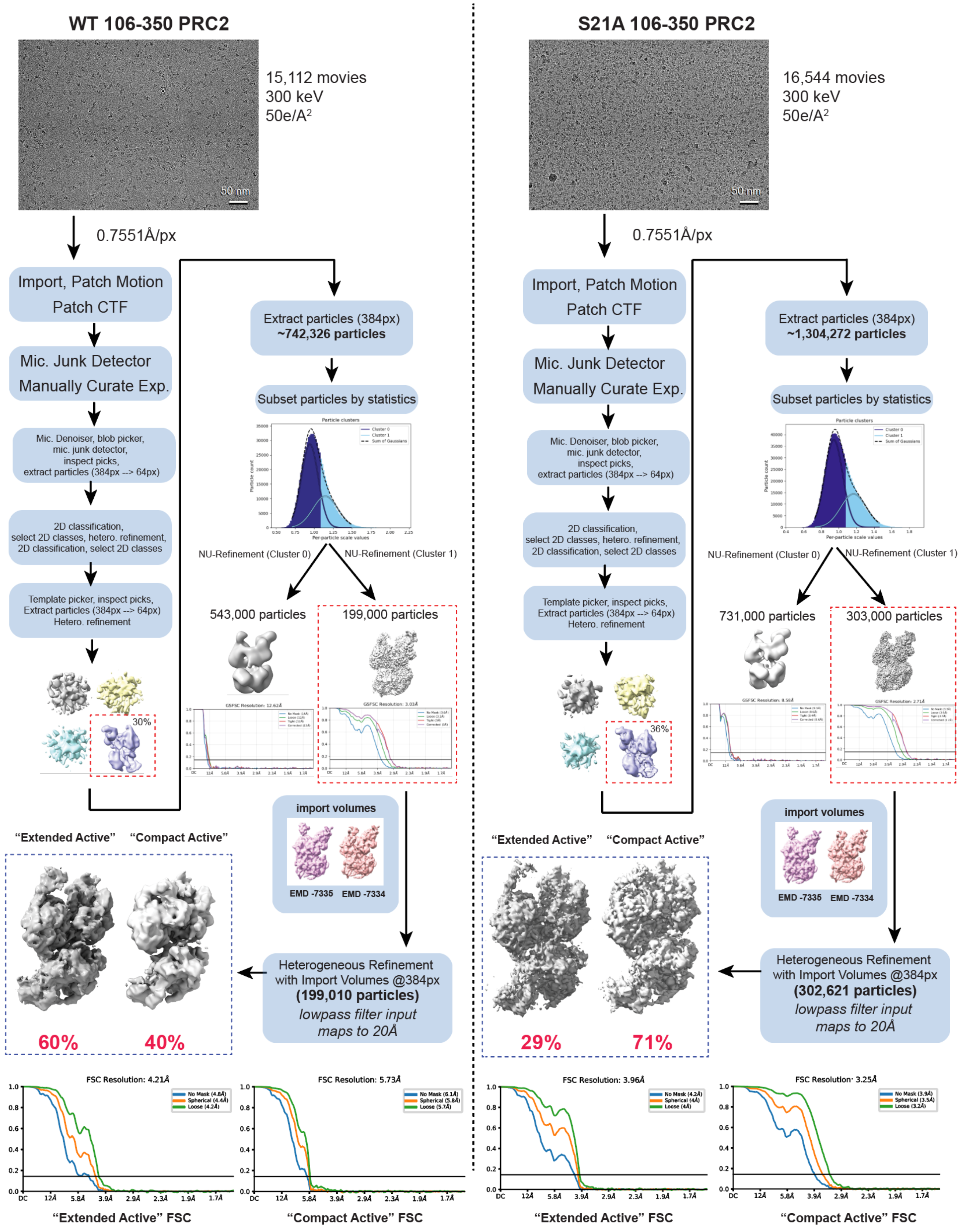
WT 106-350 PRC2
15,112 movies
300 keV
50e/A2
50 nm
0.7551Å/px
Import, Patch Motion
Patch CTF
Mic. Junk Detector
Manually Curate Exp.
Mic. Denoiser, blob picker, mic. junk detector, inspect picks, extract particles (384px --> 64px)
2D classification, select 2D classes, hetero. refinement, 2D classification, select 2D classes
Template picker, inspect picks, Extract particles (384px --> 64px) Hetero. refinement
30%
Extract particles (384px)
~742,326 particles
Subset particles by statistics
NU-Refinement (Cluster 0)
NU-Refinement (Cluster 1)
543,000 particles
199,000 particles
import volumes
EMD -7335
EMD -7334
"Extended Active"
"Compact Active"
60%
40%
Heterogeneous Refinement with Import Volumes @384px
(199,010 particles)
lowpass filter input maps to 20Å
FSC Resolution: 4.21Å
FSC Resolution: 5.73Å
"Extended Active" FSC
"Compact Active" FSC
S21A 106-350 PRC2
16,544 movies
300 keV
50e/A2
50 nm
0.7551Å/px
Import, Patch Motion
Patch CTF
Mic. Junk Detector
Manually Curate Exp.
Mic. Denoiser, blob picker, mic. junk detector, inspect picks, extract particles (384px --> 64px)
2D classification, select 2D classes, hetero. refinement, 2D classification, select 2D classes
Template picker, inspect picks, Extract particles (384px --> 64px) Hetero. refinement
36%
Extract particles (384px)
~1,304,272 particles
Subset particles by statistics
NU-Refinement (Cluster 0)
NU-Refinement (Cluster 1)
731,000 particles
303,000 particles
import volumes
EMD -7335
EMD -7334
"Extended Active"
"Compact Active"
29%
71%
Heterogeneous Refinement with Import Volumes @384px
(302,621 particles)
lowpass filter input maps to 20Å
FSC Resolution: 3.96Å
FSC Resolution: 3.25Å
"Extended Active" FSC
"Compact Active" FSC

**Figure S2. Cryo-EM data-processing workflows for WT and EZH2 S21A PRC2-AEBP2-JARID2(106-350) complexes, related to Figure 2.**

WT and EZH2 S21A datasets were collected in parallel and processed using the same workflow. Representative micrographs are shown with 50-nm scale bars. Data were collected at 300 keV with a total electron exposure of 50 e−/Å² and a physical pixel size of 0.7551 Å/pixel. Preprocessing was performed in cryoSPARC v4.7.1. Following patch motion correction, patch CTF estimation, micrograph screening, and manual curation, particles were identified using micrograph denoising, blob picking, and junk detection. Particles were initially extracted at 64 pixels and subjected to 2D classification and heterogeneous refinement. Selected classes were used for template picking, after which particles were re-extracted at 384 pixels, subset by particle statistics, and subjected to non-uniform refinement. Cluster-derived particle subsets were then analyzed by heterogeneous refinement using 20 Å low-pass-filtered imported volumes corresponding to the extended-active and compact-active states (EMD-7335 and EMD-7334). The resulting particle distributions and FSC curves are shown for WT and EZH2 S21A PRC2.

**Figure S3**

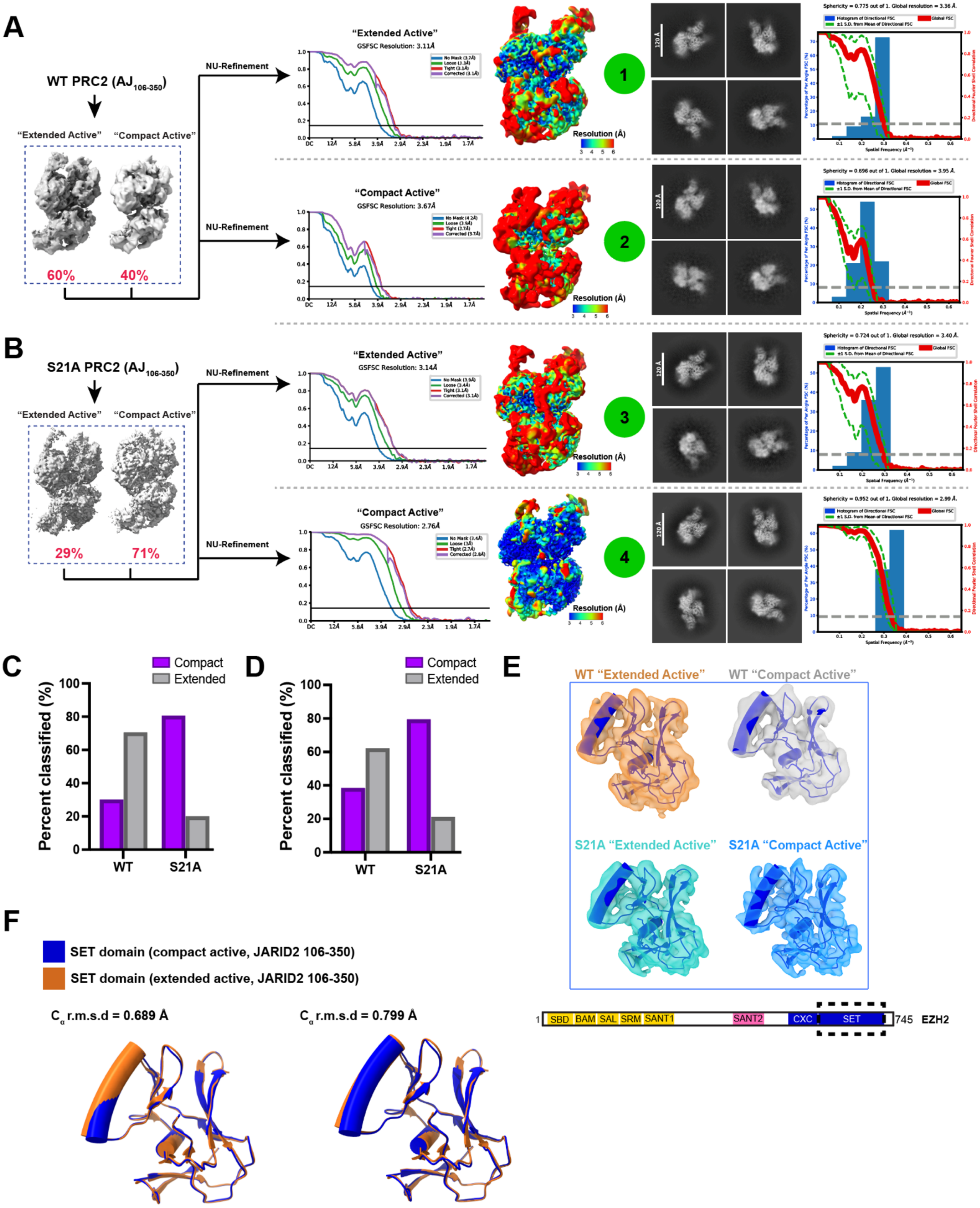

A
WT PRC2 (AJ106-350)
"Extended Active"
"Compact Active"
60%
40%
NU-Refinement
"Extended Active"
GSFSC Resolution: 3.11Å
"Compact Active"
GSFSC Resolution: 3.67Å
Resolution (Å)
1
2
B
S21A PRC2 (AJ106-350)
"Extended Active"
"Compact Active"
29%
71%
"Extended Active"
GSFSC Resolution: 3.14Å
"Compact Active"
GSFSC Resolution: 2.76Å
3
4
C
Compact
Extended
Percent classified (%)
WT
S21A
D
E
WT "Extended Active"
WT "Compact Active"
S21A "Extended Active"
S21A "Compact Active"
F
SET domain (compact active, JARID2 106-350)
SET domain (extended active, JARID2 106-350)
Cα r.m.s.d = 0.689 Å
Cα r.m.s.d = 0.799 Å
1 SBD BAM SAL SRM SANT1 SANT2 CXC SET 745 EZH2

**Figure S3. Cryo-EM reconstruction validation and SET-domain comparisons for WT and EZH2 S21A PRC2, related to Figure 2.**

**A.** Activator-bound wild-type PRC2 particles containing AEBP2 and JARID2(106-350) were classified into extended-active and compact-active conformational states and independently subjected to non-uniform refinement. The percentage of particles assigned to each class is indicated. Gold-standard Fourier shell correlation (GSFSC) curves, local-resolution maps, representative reference-free 2D class averages, and three-dimensional Fourier shell correlation (3DFSC) analyses are shown for each refined particle subset. Local-resolution maps are colored by resolution from 3 to 6 Å. The 3DFSC plots show directional FSC distributions, global FSC curves, map sphericity, and global resolution for each reconstruction.

**B.** Activator-bound EZH2 S21A PRC2 particles containing AEBP2 and JARID2(106-350) were classified into extended-active and compact-active conformational states and independently subjected to non-uniform refinement. The percentage of particles assigned to each class is indicated. GSFSC curves, local-resolution maps, representative reference-free 2D class averages, and 3DFSC analyses are shown for each refined particle subset. Local-resolution maps are colored by resolution from 3 to 6 Å. The 3DFSC plots show directional FSC distributions, global FSC curves, map sphericity, and global resolution for each reconstruction.

**C.** Particle-distribution bar plots for WT and EZH2 S21A particles from heterogeneous refinements performed with input maps low-pass filtered to 30 Å and forced hard classification enabled. Particle distributions were WT compact, 29.8%; WT extended, 70.2%; S21A compact, 80.3%; and S21A extended, 19.7%.

**D.** Particle-distribution bar plots for WT and EZH2 S21A particles from heterogeneous refinements performed with input maps low-pass filtered to 20 Å and forced hard classification enabled. Particle distributions were WT compact, 38.2%; WT extended, 61.8%; S21A compact, 79.2%; and S21A extended, 20.8%.

**E.** Extracted cryo-EM densities and rigid-body fitting of the EZH2 SET domain in WT and EZH2 S21A PRC2 extended-active and compact-active states. Previously determined EZH2 SET-domain models from compact-active PRC2 (PDB 6C23) and extended-active PRC2 (PDB 6C24) were fit into the corresponding experimental maps. The EZH2 domain schematic indicates the SET domain analyzed.

**F.** Structural comparison of the EZH2 SET domain in compact-active and extended-active PRC2-AEBP2-JARID2(106-350) conformations (PDB 6C23 and 6C24). Compact-active and extended-active SET domains are shown in blue and orange, respectively. Left, alignment based on the SET domain. Right, comparison after alignment based on EED. Cα r.m.s.d. values are indicated.

### Figure S4

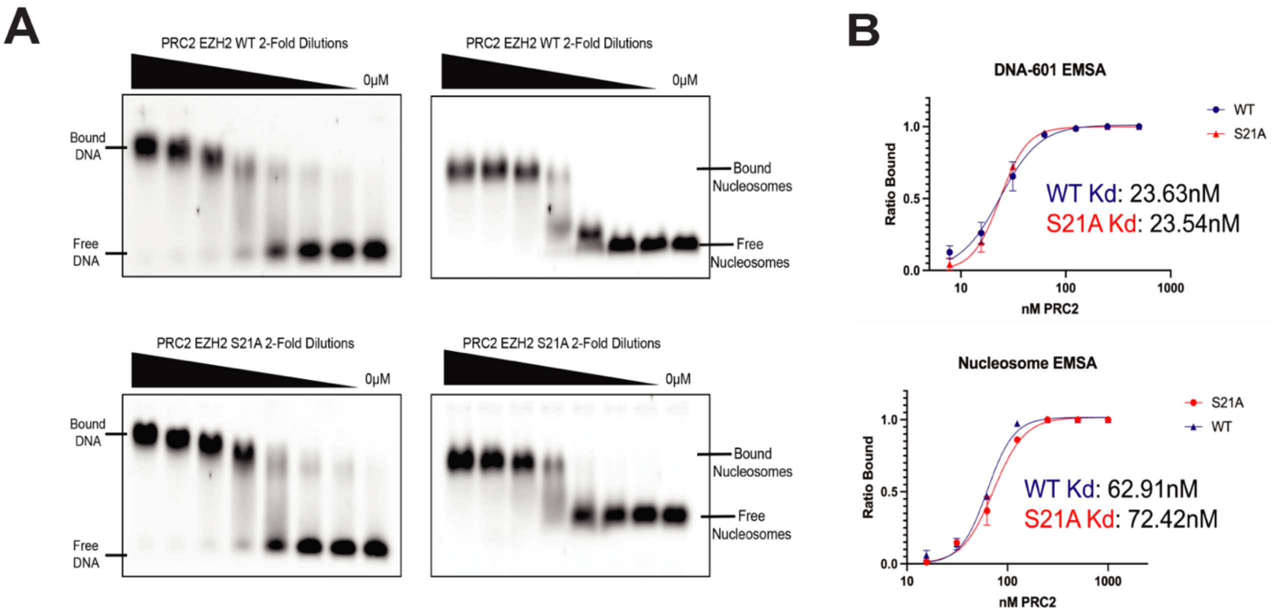


## C

| H3K27me3 (μM) | Enzyme | kcat (min⁻¹) | Km (μM) | kcat/Km (μM⁻¹·min⁻¹) |
|---|---|---|---|---|
| 0 μM | WT | 18.6 ± 1.2 | 114.0 ± 14.8 | 0.2 ± 0.0 |
| | S21A | 22.5 ± 1.1 | 58.8 ± 6.9 | 0.4 ± 0.1 |
| 10 μM | WT | 37.8 ± 2.5 | 43.8 ± 8.0 | 0.9 ± 0.2 |
| | S21A | 55.2 ± 3.3 | 23.6 ± 4.3 | 2.3 ± 0.5 |
| 25 μM | WT | 98.4 ± 7.7 | 54.5 ± 9.5 | 1.8 ± 0.4 |
| | S21A | 98.1 ± 5.1 | 32.7 ± 4.5 | 3.0 ± 0.4 |
| 40 μM | WT | 75.2 ± 2.8 | 18.6 ± 2.4 | 4.1 ± 0.6 |
| | S21A | 88.8 ± 4.9 | 17.9 ± 3.5 | 4.9 ± 1.0 |

**Figure S4. DNA and nucleosome binding and kinetic characterization of WT and EZH2 S21A PRC2, related to Figure 3.**

**A.** Representative electrophoretic mobility shift assay (EMSA) gels measuring binding of WT or EZH2 S21A PRC2 to Cy5-labeled 601 DNA and Cy5-labeled 601 nucleosomes. PRC2 was titrated in 2-fold serial dilutions beginning at 500 nM for DNA and 1,000 nM for nucleosomes, with a no-PRC2 control indicated.

**B.** Quantification of DNA- and nucleosome-binding EMSAs. Bound fraction is plotted as a function of PRC2 concentration, and fitted curves were used to derive apparent dissociation constants. Points show mean ± SD from independent experiments.

**C.** Summary of Michaelis-Menten kinetic parameters for recombinant PRC2 containing WT EZH2 or EZH2 S21A assayed with histone H3(18-37) peptide substrate in the presence of 0, 10, 25, or 40 μM stimulatory H3K27me3 peptide. Values for $k_{cat}$, $K_m$, and $k_{cat}/K_m$ are reported as fitted estimates ± standard errors and were derived from the kinetic analyses shown in **Figure 3B**.

### Figure S5

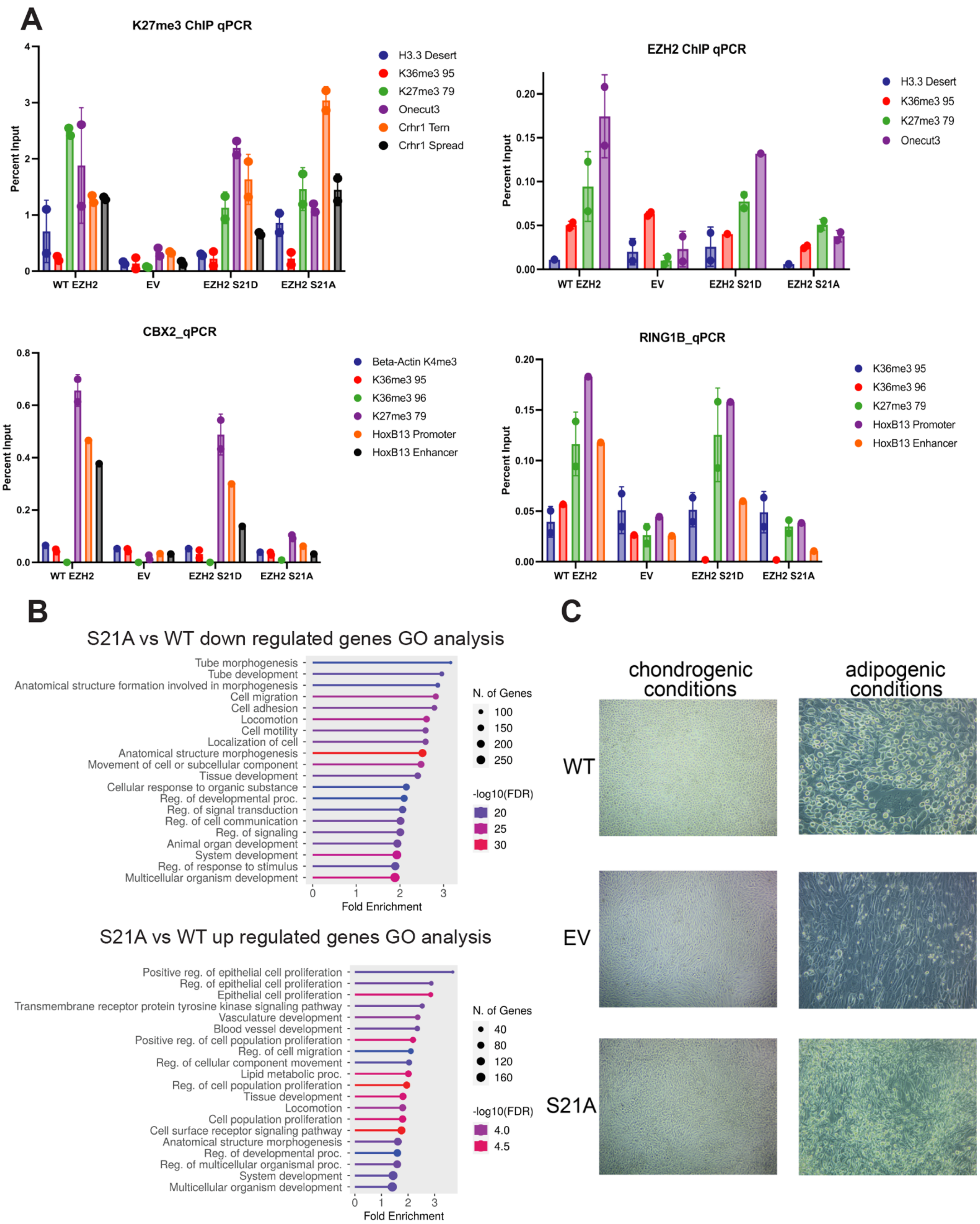
A
K27me3 ChIP qPCR
Percent Input
WT EZH2
EV
EZH2 S21D
EZH2 S21A
H3.3 Desert
K36me3 95
K27me3 79
Onecut3
Crhr1 Tern
Crhr1 Spread
EZH2 ChIP qPCR
CBX2_qPCR
Beta-Actin K4me3
K36me3 96
HoxB13 Promoter
HoxB13 Enhancer
RING1B_qPCR
B
S21A vs WT down regulated genes GO analysis
Tube morphogenesis
Tube development
Anatomical structure formation involved in morphogenesis
Cell migration
Cell adhesion
Locomotion
Cell motility
Localization of cell
Anatomical structure morphogenesis
Movement of cell or subcellular component
Tissue development
Cellular response to organic substance
Reg. of developmental proc.
Reg. of signal transduction
Reg. of cell communication
Reg. of signaling
Animal organ development
System development
Reg. of response to stimulus
Multicellular organism development
Fold Enrichment
N. of Genes
-log10(FDR)
S21A vs WT up regulated genes GO analysis
Positive reg. of epithelial cell proliferation
Reg. of epithelial cell proliferation
Epithelial cell proliferation
Transmembrane receptor protein tyrosine kinase signaling pathway
Vasculature development
Blood vessel development
Positive reg. of cell population proliferation
Reg. of cell migration
Reg. of cellular component movement
Lipid metabolic proc.
Reg. of cell population proliferation
Tissue development
Locomotion
Cell population proliferation
Cell surface receptor signaling pathway
Anatomical structure morphogenesis
Reg. of developmental proc.
Reg. of multicellular organismal proc.
System development
Multicellular organism development
C
chondrogenic conditions
adipogenic conditions
WT
EV
S21A

**Figure S5. Chromatin occupancy, gene ontology, and early mesenchymal differentiation phenotypes, related to Figures 4 and 5.**

**A.** ChIP-qPCR analysis of H3K27me3, EZH2, CBX2, and RING1B enrichment at representative Polycomb target loci and negative-control regions. Enrichment is shown for C3H10T1/2 *Ezh2*-knockout mesenchymal cells expressing wild-type EZH2, empty vector, S21D, or S21A, permitting comparison of PRC2 activity and occupancy with canonical PRC1 occupancy.

**B.** Gene ontology (GO) enrichment analysis of downregulated (top) and upregulated (bottom) genes in EZH2 S21A-expressing mesenchymal cells relative to wild-type EZH2-expressing cells. Dot size indicates the number of genes, color indicates $-\log_{10}$(FDR), and the x-axis shows fold enrichment.

**C.** Representative images of *Ezh2*-knockout C3H10T1/2 mesenchymal progenitor cells expressing WT EZH2, empty vector, or EZH2 S21A after 5 days under chondrogenic (left) or adipogenic (right) differentiation conditions. Cultures were fixed and imaged at day 5 to visualize early morphological changes associated with chondrocyte and adipocyte lineage commitment.
